# Pathological and metabolic underpinnings of energetic inefficiency in temporal lobe epilepsy

**DOI:** 10.1101/2021.09.23.461495

**Authors:** Xiaosong He, Lorenzo Caciagli, Linden Parkes, Jennifer Stiso, Teresa M. Karrer, Jason Z. Kim, Zhixin Lu, Tommaso Menara, Fabio Pasqualetti, Michael R. Sperling, Joseph I. Tracy, Dani S. Bassett

**Author notes:** Correspondence to the below authors: Dr. Xiaosong He, Dr. Dani S. Bassett.

## Abstract

The human brain consumes a disproportionate amount of energy to generate neural dynamics. Yet precisely how energetic processes are altered in neurological disorders remains far from understood. Here, we use network control theory to profile the brain’s energy landscape, describing the rich dynamical repertoire supported by the structural connectome. This approach allows us to estimate the energy required to activate a circuit, and determine which regions most support that activation. Focusing on temporal lobe epilepsy (TLE), we show that patients require more control energy to activate the limbic network than healthy volunteers, especially ipsilateral to the seizure focus. Further, greater energetic costs are largely localized to the ipsilateral temporo-limbic regions. Importantly, the energetic imbalance between ipsilateral and contralateral temporo-limbic regions is tracked by asymmetric metabolic patterns, which in turn are explained by asymmetric gray matter volume loss. In TLE, failure to meet the extra energy demands may lead to suboptimal brain dynamics and inadequate activation. Broadly, our investigation provides a theoretical framework unifying gray matter integrity, local metabolism, and energetic generation of neural dynamics.

## Main

Human brain function emerges from continuous neural dynamics that give rise to diverse activation states and rich rules for transitioning between states^1–5^. Those dynamics pose marked energetic demands, consuming 20% of the body’s energy while comprising merely 2% of the body’s weight^6^. Pathological disruptions to healthy neural dynamics and their energetic sequelae, as observed in neurological and psychiatric diseases, can have devastating consequences for cognitive function and behavior. One prototypical example is epileptic seizures, which are transient periods of pathological hypersynchronous neuronal activities^7^. Seizures consume significant energy and instantly disrupt ongoing brain dynamics, causing marked behavioral disturbances^8,9^. Despite their transience, the impact of seizures on brain function can linger well beyond seizure termination, especially in patients with drug-resistant focal epilepsy, such as temporal lobe epilepsy (TLE). Neuropsychological assessments performed during the interictal period, *i*.*e*., when patients are not suffering from seizures, reveal persistent deficits in multiple cognitive and affective domains^10–13^, indicating the presence of prolonged disruptions to normal brain dynamics. Little is known about this pervasive neural dysfunction throughout the vast temporal epochs between seizures, its relation to brain energetics, and its potential dependence upon the brain’s structural backbone.

TLE is marked by widespread alterations in brain structure, which—importantly—extends beyond the seizure foci^14–16^. Concordantly, epileptogenic regions evince persistent hypometabolism during interictal periods^17^. The severity of both structural damage and hypometabolism is associated with cognitive decline in TLE patients^18–20^, suggesting that chronic neural dysfunction might be rooted in a reduced baseline metabolism underpinned by compromised structural integrity. Regional hypometabolism could hamper the attainment and maintenance of healthy activation levels, in turn decrementing the brain’s dynamic repertoire and clamping cognitive function. Examples of such altered dynamics in TLE abound, spanning reduced language network flexibility (*i*.*e*., fewer state transitions)^21^, reduced memory network flexibility^22^, delayed information flow, and slower activation spreading times^23^. Despite these pervasive alterations in dynamics, metabolism, and structure, little is known about the mechanistic relationships between them.

To formally assess how damage to structural connectivity disrupts energetic generation of brain dynamics, we use network control theory (NCT), a powerful approach from systems engineering typically deployed to design and manage technological, robotic, and communication systems. NCT allows us to evaluate the energetic cost of brain states—and transitions between them—as a function of the underlying network architecture. We begin by stipulating a dynamical model whereby activity is constrained to spread along structural connections (**Figure 1a**). Using this model, we can quantify the control energy required to move between any two states or patterns of activity^24^. Prior work has shown that the control energy required to transition between states varies with cognitive demand^25^, decreases over development^26^, and is altered in psychiatric disorders^27,28^. Prior studies have also demonstrated that control energy can be causally manipulated by brain stimulation^29^ and antipsychotic medication^25^. These efforts lay important groundwork for our use of NCT to model the structurally-constrained energetic processes of brain state transitions in patients with TLE, by supporting its feasibility, ensuring its methodological rigor, and underscoring its biological relevance.

**Figure 1.**
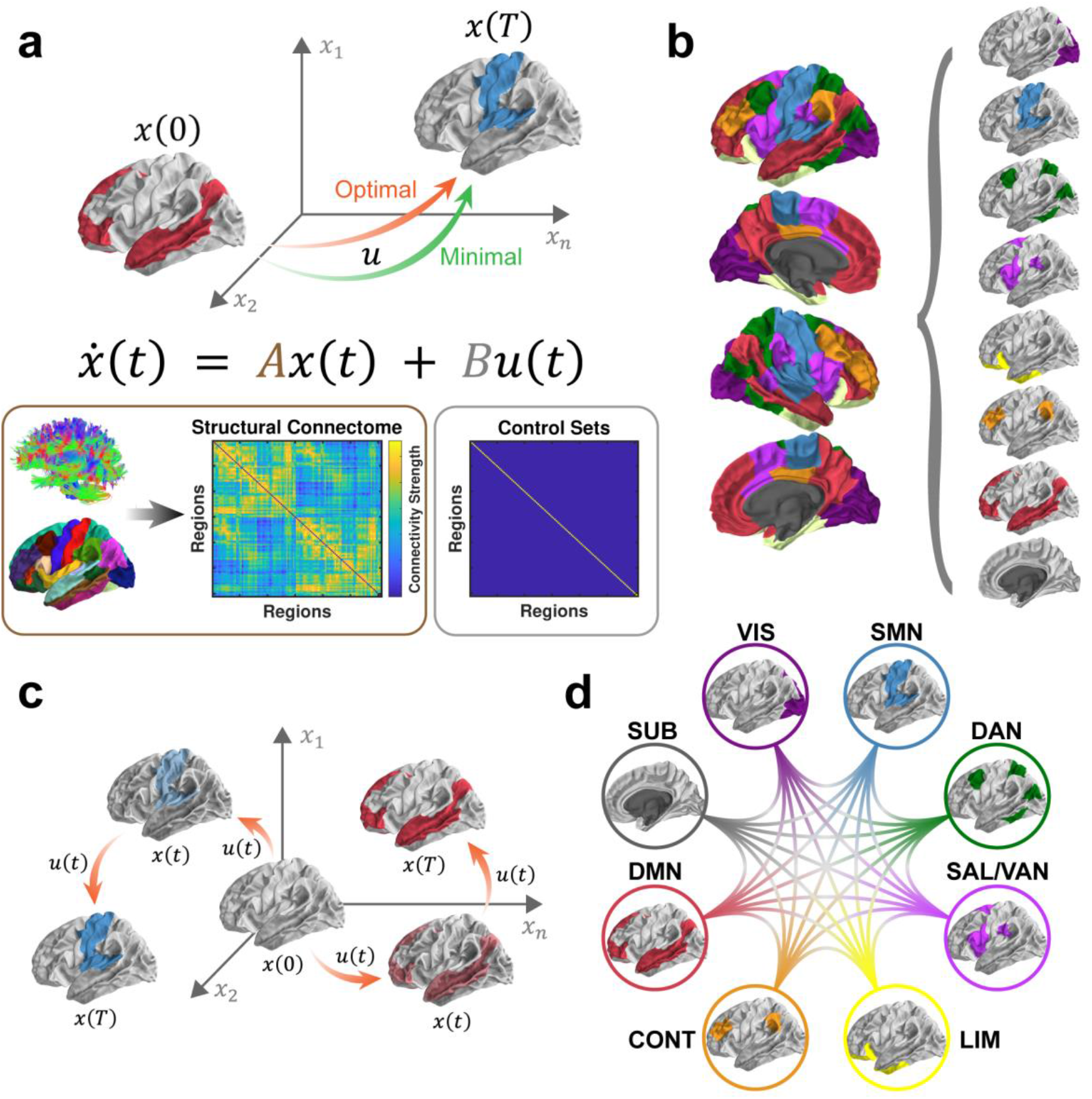
Schematic of methods. (a) Based on a simplified noise-free, linear, continuous-time, and time-invariant model of neural dynamics, we simulate energetic processes during brain state transitions instantiated upon and constrained by the structural connectome (matrix A). Two types of control energy (a quadratic function of *u*) are depicted: the minimum control energy required to drive the brain from an initial state [*x(0)*] to a final state [*x(T)*] using a specific set of control nodes (whole brain, matrix B); the optimal control energy additionally constrains the length of the trajectory between states. (b) Eight preferential brain states are defined according to the known intrinsic connectivity networks (ICN)^32,33^. Within each state, regions from a specific ICN are activated at a magnitude of 1, whereas the rest of the brain remains at 0 (inactivated). These preferential brain states constitute the initial and final states of our simulations. (c) We then simulate the energetic inputs required to activate each of the preferential brain states from a theoretical baseline (*i*.*e*., activity magnitude of 0). Next, we estimate the optimal control energy consumed during each of the activation processes across the whole brain for each subject. (d) We also simulate transitions between preferential brain states, and estimate the minimal control energy consumed at each brain region for each subject. Abbreviations: VIS, visual network; SMN, somatomotor network; DAN, dorsal attention network; SAL/VAN, salience/ventral attention network; CONT, executive control network; DMN, default mode network; SUB, subcortical network.

Although a generic artificial system could hypothetically visit any activation state, evidence suggests that the brain preferentially visits some states more often than others^4,30,31^. These preferential states can be defined by the co-activation of regions that are functionally coupled at rest^32,33^. Such so-called *intrinsic connectivity networks* (ICNs) can also be detected during task performance, and have been shown to support a range of cognitive processes^32,34,35^. Here, we study the efficient attainment of eight such preferential states^33,36^, whereby only regions in a given ICN are active (**Figure 1b**). Then, we use NCT to simulate transitions among preferential states and to estimate the associated energy costs, thereby probing the brain’s efficiency and integrity in the presence of TLE. Our simulations evaluate two transition types: (1) *reaching transitions*, where the brain moves from a theoretical baseline to a preferential state (**Figure 1c**); and (2) *switching transitions*, where the brain moves between two preferential states (**Figure 1d**). By estimating the control energy for *reaching transitions*, we determine which preferential states are difficult to reach; that determination informs our understanding of the ICNs impacted by TLE. Subsequently, we compute control energy for *switching transitions*, which allows us to identify the regions that tend to carry the greatest energetic burdens in supporting the brain’s dynamical repertoire.

After using NCT to unravel the energetic basis of brain dysfunction in TLE, we next dig deeper into the neurophysiological underpinnings of control energy. In the absence of any external input to the brain (*e*.*g*., brain stimulation), control energy is thought to track the cost of endogenous resources associated with internal cognitive demand^25,37^. However, there is as yet no precise evidence linking this metric to a direct readout of such resources. Here we provide exactly such evidence. As part of presurgical evaluation in patients with TLE, fluorodeoxyglucose (FDG)-PET is commonly performed interictally to measure baseline metabolic levels. Leveraging this additional piece of information, we can verify whether regions that show altered energetic efficiency in facilitating brain state transitions also present with metabolic anomalies. Moreover, we can determine how both the theoretical and empirical measures of energy costs are related to the underlying structural integrity of those regions.

In this study, we enrolled 60 TLE patients and 50 demographically matched healthy controls (HCs) (**Table 1**), who underwent an MRI scanning session including both a high angular resolution diffusion imaging (HARDI) scan and a T1-weighted (T1w) anatomical scan. Among the enrolled patients, 50 also received an FDG-PET scan as part of their presurgical evaluation. From each participant’s HARDI data, we generated a structural connectome and estimated the control energy required to perform all *reaching* and *switching transitions*. Then, we tested for energetic deficiencies in TLE by comparing control energy between patients and HCs. We showed that TLE patients present with an energetic deficiency in reaching a preferential state predominantly comprised of limbic regions. This deficiency was due to excessive energy costs associated with activating the limbic network ipsilateral to the patient’s seizure focus. When considering *switching transitions*, we found that the mesial and inferior parts of the ipsilateral temporal lobe demanded greater energy consumption in TLE than in HCs. These increased costs of regulating brain dynamics incurred by TLE patients limit their capacity to activate and maintain desired brain states, potentially leading to dysfunction. Furthermore, we found that this energetic imbalance between ipsilateral mesial and inferior temporal regions and their contralateral counterparts were accompanied by similar asymmetries in metabolic patterns, both of which are rooted in a corresponding asymmetry of underlying gray matter volume loss.

**Table 1.**
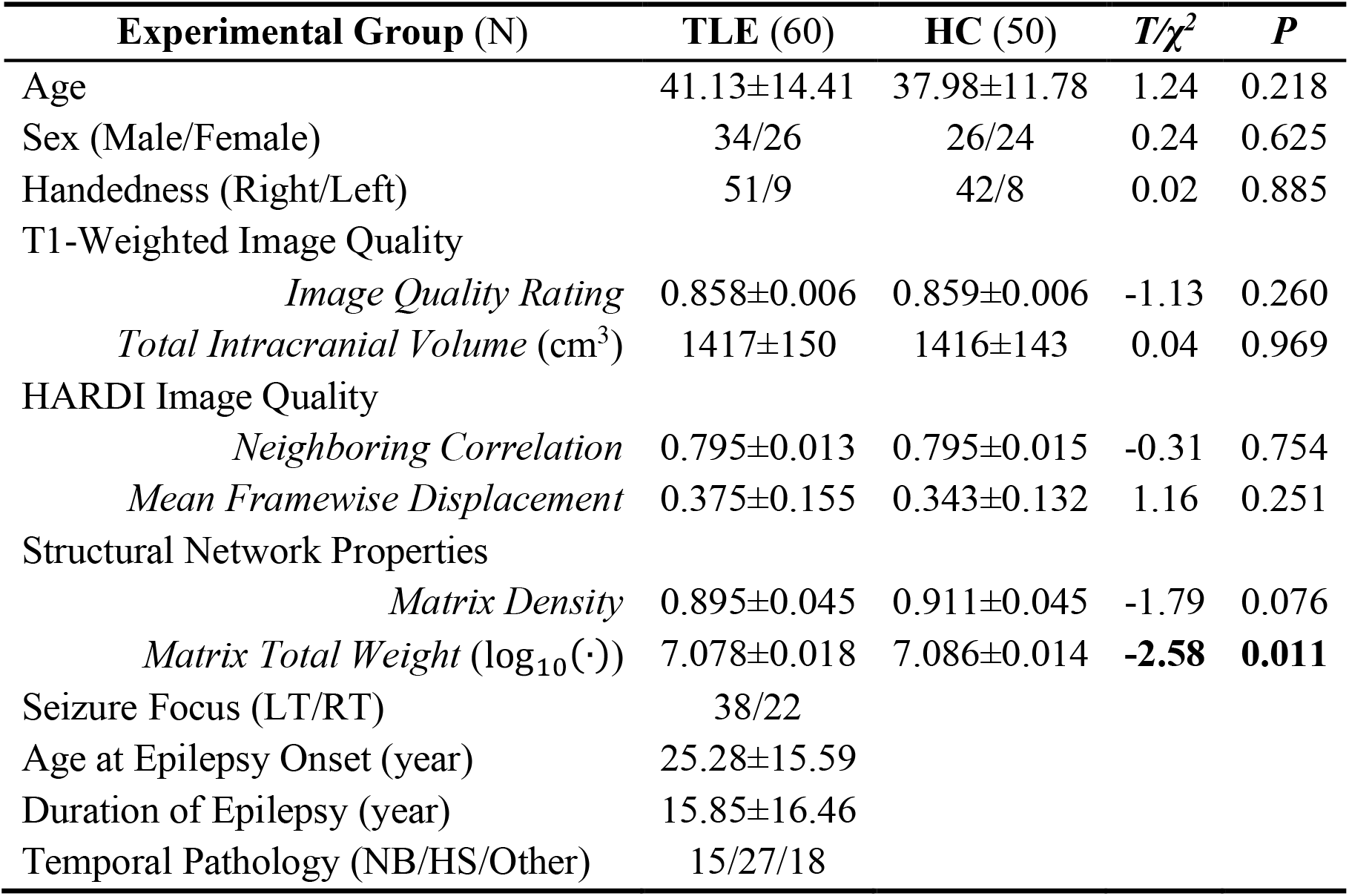
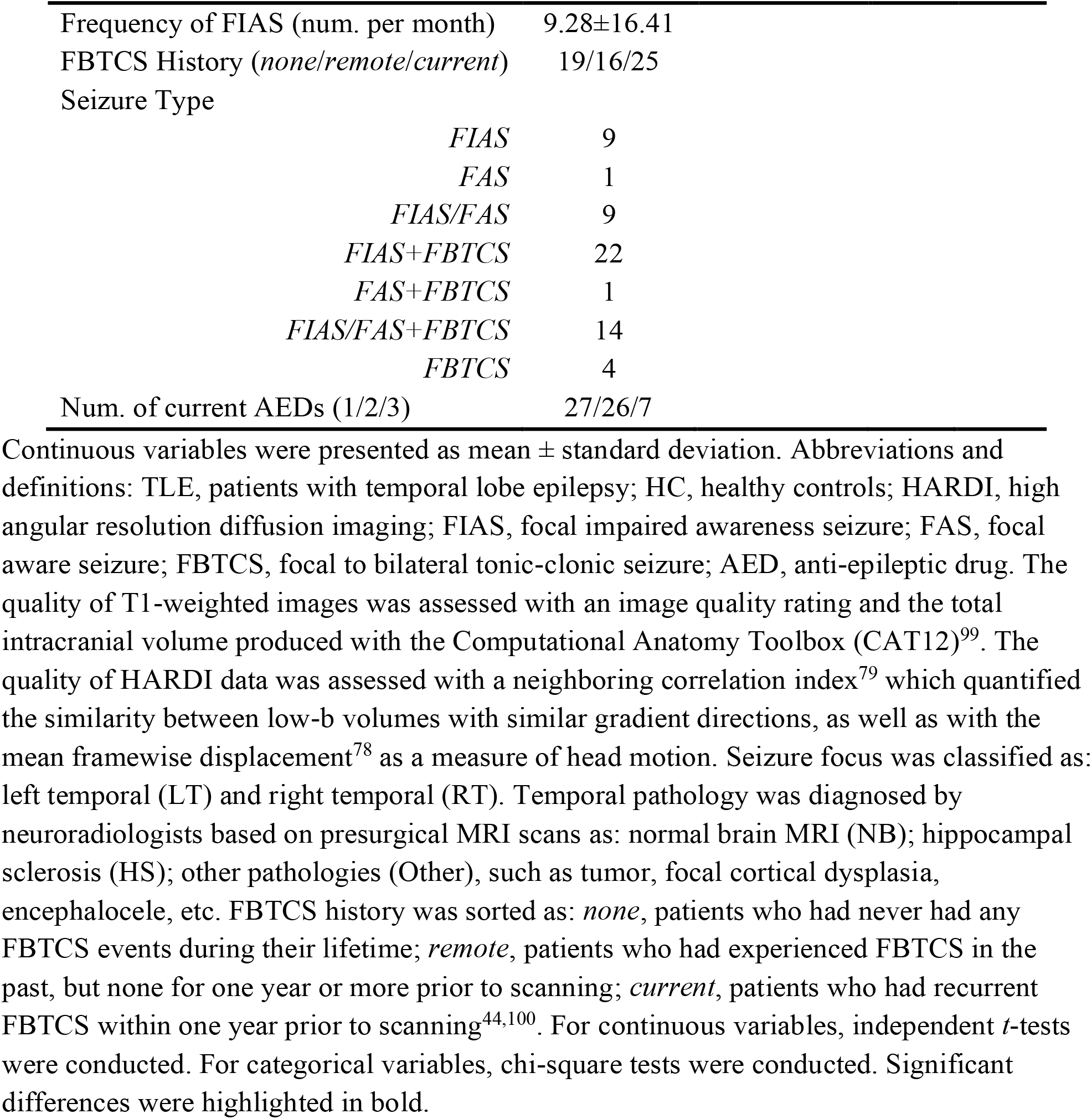
Sample demographic and clinical characteristics.

## Results

We started our analyses by verifying that the imaging data quality was comparable between the two experimental groups (**Table 1**). Next, for each participant, we reconstructed a structural white matter network as a weighted undirected adjacency matrix comprised of 122 cortical and subcortical regions (see details in *Methods*), which formed the basis of our NCT analyses (*e*.*g*., matrix A in **Figure 1a**). We observed that TLE patients presented lower matrix density and total weight (*i*.*e*., sum of all edge weights) than HCs (**Table 1**), which is in accord with the well-recognized white matter abnormalities in TLE^38,39^. To minimize the influence of demographic and data quality metrics on our subsequent statistical analyses, we regressed them out from all imaging derivates using linear regression (see *Methods*).

### Simulated activation of intrinsic connectivity networks

Our first research question pertained to the energetic costs associated with *reaching* each preferential brain state. Specifically, we examined the extent to which TLE patients exhibited energetic abnormalities during the activation of eight canonical ICNs^33,36^, including the visual, somatomotor, dorsal attention, salience/ventral attention, limbic (including amygdala and hippocampus), executive control, default mode, and subcortical networks (**Figure 1b**). We used the optimal control framework^26,29,36,40^ to model how the brain’s underlying structural network facilitated transitions from an initial baseline state to each preferential (final) state. Here, the initial state was set at a theoretical baseline with activity magnitude in all regions at 0. For each of our 8 preferential states, the activity magnitude of regions within a specific ICN was set to 1, while the rest of the brain remained at 0^26^. Thus, these *reaching transitions* simulated the rise of activity in the target ICN from a mean-centered baseline, mimicking the activation process triggered by specific cognition operations. For this model, an optimal solution of the control energy needed at each region can be produced by constraining both the energy costs and the length of the transition trajectory based on the underlying network topology^37^. As previously^37^, we referred to this solution as the *optimal control energy* (OCE), and summarized it globally as a measure of the brain’s energetic efficiency (**Figure 1c**, see details in *Methods*). For each participant, we estimated the global OCE during each *reaching transition*. We regressed the confounding variables out of these global OCE estimates and compared the residuals between our two groups using a permutation-based *t*-test^41^, an approach that simultaneously corrects for multiple comparisons by controlling the family-wise error rate^42^. We found that TLE patients required greater global OCE to activate the limbic network compared to HCs (Welch’s *t*_108_=3.80, *P*_corr_=0.002). The global OCE needed to activate other ICNs did not significantly differ between the two groups (Welch’s |*t*_108_’s|<1.55, *P*_corr_>0.609) (**Figure 2**). This finding suggests that it is energetically more challenging for TLE patients to specifically activate the limbic network.

**Figure 2.**
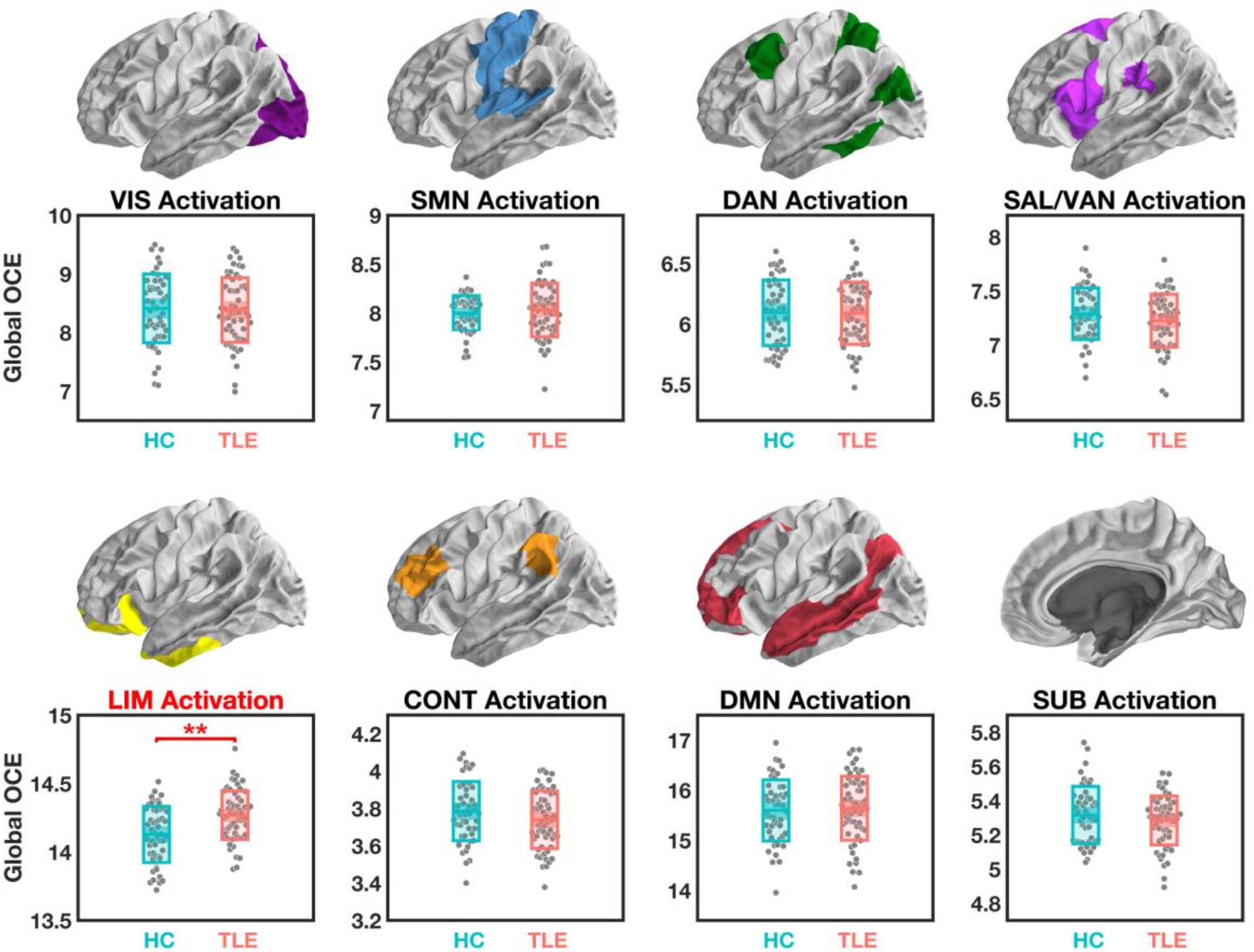
Global optimal control energy (OCE) estimated during simulated activation of intrinsic connectivity networks. After correction for multiple comparisons, significant group differences were only found for the simulated activation of the limbic network (LIM), which demanded more global OCE in patients with temporal lobe epilepsy (TLE) compared to healthy controls (HC). Other abbreviations: VIS, visual network; SMN, somatomotor network; DAN, dorsal attention network; SAL/VAN, salience/ventral attention network; CONT, executive control network; DMN, default mode network; SUB, subcortical network. ^**^, *P*_corr_ < 0.01. The central mark indicates the median, and the bottom and top edges of the box indicate the 25^th^ and 75^th^ percentiles, respectively.

In the above analysis, our preferential states were defined with ICNs extended into both hemispheres. However, seizures in our enrolled TLE patients were exclusively of unilateral origin, *i*.*e*., from either the left or the right temporal lobe. This laterality of seizure focus suggests that the energetic inefficiency we observed in these patients may be asymmetric, especially with regard to the limbic network, which includes both the epileptogenic temporal lobe and its contralateral counterpart. To probe this asymmetry, we re-simulated the *reaching transition* for the limbic network twice, once to activate limbic regions in the left hemisphere only, and once to activate regions in the right hemisphere. Although such a hemispheric restriction of activation is unlikely to occur in the brain, the simulation offers an opportunity to assess the laterality of the pathological burden observed in TLE patients. When examining lateralized global OCE, we found a significant hemisphere-by-group interaction [*F*_(2,107)_=15.20, *P*=2×10^−6^] (**Figure 3**), demonstrating that TLE patients required more energy to activate the limbic network ipsilateral to the seizure focus. More specifically, TLE patients with a left-sided seizure focus required more energy to activate the left-hemispheric limbic network (vs. right TLE: *P*_Bonferroni_=0.048; vs. HC: *P*_Bonferroni_=4×10^−5^), whereas TLE patients with a right-sided seizure focus required more energy to activate the right-hemispheric limbic network (vs. left TLE (*P*_Bonferroni_=0.007); vs. HC: *P*_Bonferroni_=2×10^−4^). By contrast, we observed no significant differences between TLE patients and HCs in the energy needed to activate the contralateral limbic network (*P*_Bonferroni_’s>0.588). Thus, the extra energetic costs associated with limbic network activation in TLE can be attributed to the increased energetic needs of the ipsilateral hemisphere, but not of the contralateral one.

**Figure 3.**
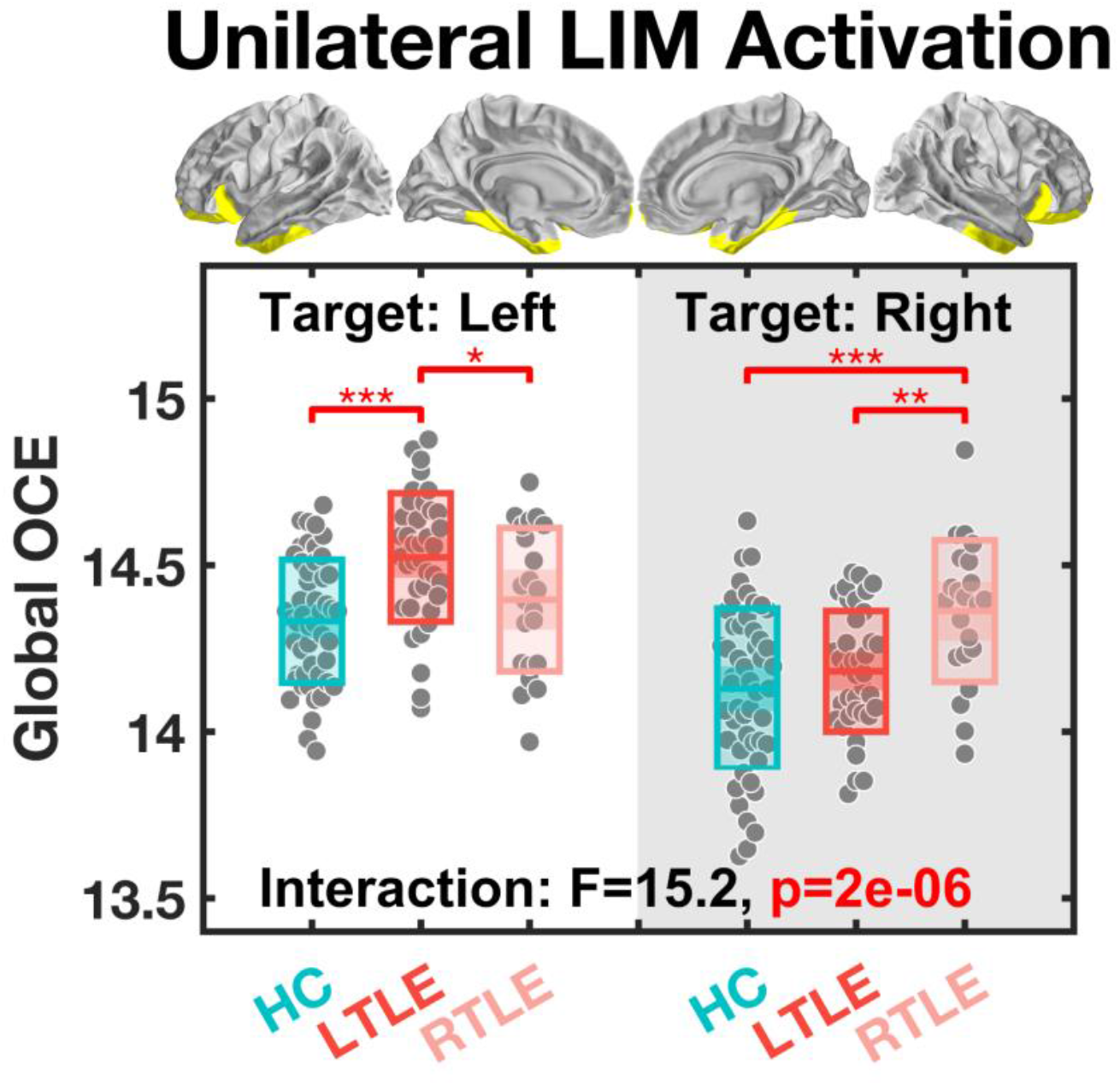
Global optimal control energy (OCE) estimated during a simulated transition from the baseline to a final state where only one side of the limbic network (LIM) is activated. When the target was set to the left hemispheric LIM, only left temporal lobe epilepsy patients (LTLE) needed more energy than the other two groups. When the target was set to the right hemispheric LIM, only right TLE patients (RTLE) needed more energy than the other two groups. Other abbreviations: HC, healthy controls. ^*^, *P*_Bonferroni_<0.05, ^**^, *P*_Bonferroni_<0.01, ^***^, *P*_Bonferroni_<0.001. The central mark indicates the median, and the bottom and top edges of the box indicate the 25^th^ and 75^th^ percentiles, respectively.

### Regional energetic efficiency in supporting brain state transitions

Empirical brain dynamics depend not only on the attainment of different states, but even more on the flexible transitions among them, supported by brain regions consuming energy at different efficiencies. While our *reaching transition* simulations enabled us to identify global energetic inefficiency during limbic network activation in TLE patients, such transitions are not necessarily representative enough, *i*.*e*., the brain does not revisit a specific baseline each time, but rather, continuously transitions between different states. Thus, to better profile regional energetic efficiency, we extended our investigation to examine *switching transitions* among our preferential states, as a closer approximation of empirical brain dynamics^36^. For each subject, we modeled a total of 64 pairwise transitions, including both reciprocal state transitions and single state persistence (*i*.*e*., transition starts and ends at the same state) among the eight ICN-defined preferential states (**Figure 1d**). To allow maximal flexibility during the simulated transitions, we estimate the *minimal control energy* (MCE), which can be obtained by only constraining for the control input to facilitate the designated transition regardless of its trajectory^37^. For each region, we averaged the MCE across all 64 transitions as a region-specific metric of energetic efficiency at the individual level. To enhance statistical power^38,43,44^, we mirror-flipped the regional MCE of the right TLE patients to group metrics according to the laterality of the seizure focus^38,44,45^. This procedure was done after confound regression Z-standardization of each patient’s regional MCE relative to HC data. Thus, instead of comparing raw values of regional MCE from mixed hemispheric origin between patients and HCs, we performed a permutation-based one-sample *t*-test on patients’ Z-scores of the 122 regions, to identify abnormal regional energetic efficiencies within ipsilateral and contralateral hemispheres, respectively (see *Methods* for further details).

After applying a multiple comparison correction, we found that regional MCE was significantly elevated in TLE patients within the ipsilateral hemisphere only. This elevation occurred specifically in regions such as the temporal pole (*t*_59_=5.40, *P*_corr_=10^−4^), inferior temporal gyrus (*t*_59_=5.03, *P*_corr_=5×10^−4^), amygdala (*t*_59_=6.01, *P*_corr_=10^−5^), hippocampus (*t*_59_=5.24, *P*_corr_=2×10^−4^), parahippocampal gyrus (*t*_59_=4.54, *P*_corr_=0.003), and fusiform gyrus (*t*_59_=4.93, *P*_corr_=7×10^−4^), as well as the isthmus of the cingulate gyrus (*t*_59_=4.17, *P*_corr_=0.011, rest of the brain: |*t*_59_’s|<3.41, *P*_corr_>0.114) (**Figure 4a**). No significant effects were observed in the contralateral hemisphere. In TLE, these regions required more energy to facilitate the same brain state transitions than in HC. Furthermore, this energetic inefficiency was largely located in the ipsilateral temporo-limbic regions, consistent with our previous results showing the costly activation of the limbic network in TLE patients.

**Figure 4.**
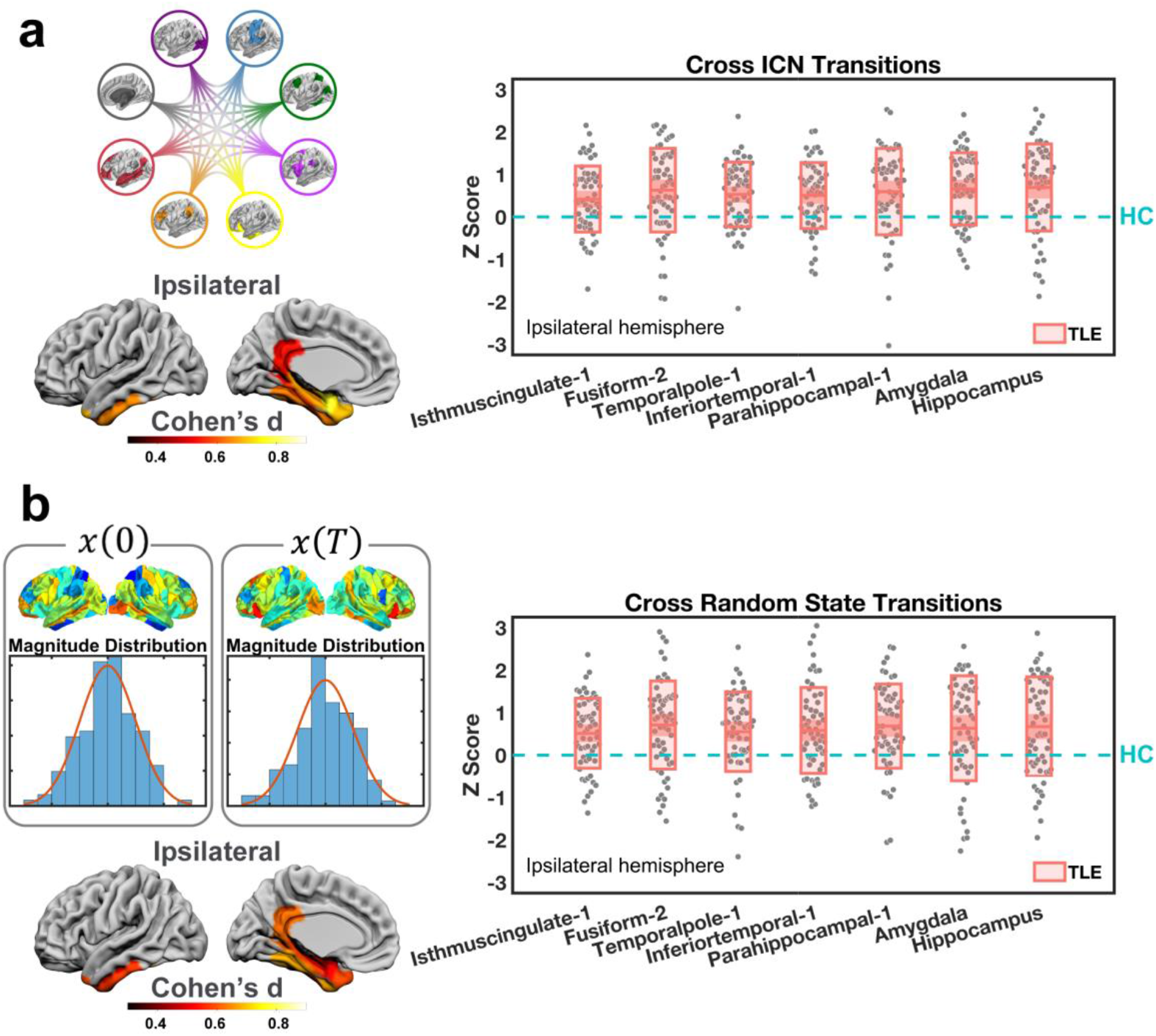
Regional energy efficiency differences between temporal lobe epilepsy (TLE) patients and healthy controls (HC). (a) We estimated the minimal control energy consumption of each region during all transitions between the brain states defined by intrinsic connectivity networks (ICNs). In the hemisphere ipsilateral to the seizure focus, we found significantly higher energy consumption in TLE patients than in HC among several temporo-limbic regions. (b) We then estimated the minimal control energy consumption of each region during transitions between 100,000 pairs of initial [*x(0)*] and final states [*x(T)*] with randomly generated activity magnitudes. Concordant results were found, showing that the patients needed significantly higher control energy in the ipsilateral temporo-limbic regions. The box plots depict the deviation scores (Z) of energy consumption of TLE patients in reference to HC. Only regions with significant group differences after multiple comparison corrections are displayed, including the isthmus of the cingulate gyrus (Isthmuscingulate-1), fusiform (Fusiform-2), temporal pole (Temporalpole-1), inferior temporal gyrus (Inferiortemporal-1), parahippocampal gyrus (Parahippocampal-1), amygdala, and hippocampus.

It remains to be determine, however, whether simulating transitions among ICN-defined brain states can provide a representative overview of all possible brain state transitions. Thus, we stringently assessed the robustness of the above findings by comparing them to MCE values derived from transitions between 100,000 pairs of random initial and final states. These random states were generated following a Gaussian distribution of activity magnitude across the 122 regions with a mean of 1 and a standard deviation of 0.1^26^. This finite repository serves as an approximation of all possible state transitions when no *a priori* brain states are explicitly defined. We adopted the same minimal control framework as above, and obtained the Z-transformed regional energy estimates. Consistent with our primary findings, we found significantly higher MCE in the ipsilateral hemisphere only, including regions such as temporal pole (*t*_59_=4.57, *P*_corr_=0.003), inferior temporal gyrus (*t*_59_=4.45, *P*_corr_=0.004), amygdala (*t*_59_=3.96, *P*_corr_=0.022), hippocampus (*t*_59_=4.44, *P*_corr_=0.004), parahippocampal gyrus (*t*_59_=5.33, *P*_corr_=2×10^−4^), and fusiform gyrus (*t*_59_=5.30, *P*_corr_=2×10^−4^), as well as the isthmus of the cingulate gyrus (*t*_59_=4.83, *P*_corr_=0.001, rest of the brain: |*t*_59_’s|<3.30, *P*_corr_>0.156) (**Figure 4b**). Thus, these randomly generated brain states yielded results matching those observed from ICN-defined brain states, supporting the notion that our preferential states appropriately represented the repertoire of empirical brain dynamics.

### Biological validation of the brain’s energetic inefficiency in TLE

Through NCT, we have established that TLE is associated with energetic inefficiency during simulated brain dynamics, not only with respect to attaining preferential states, but also in transitioning among them. Next, we sought to validate our findings using an independent measurement of neurophysiological energy consumption, FDG-PET. FDG-PET is a common clinical investigation used for seizure focus localization, and allows probing brain metabolism *in vivo* by measuring regional glucose uptake. Here, FDG-PET was acquired in a subset of 50 TLE patients during their presurgical evaluation. In the absence of HC data as baseline, we used data from the contralateral hemisphere in the same patient as a reference. After confound regression, we generated a laterality index (LI) of glucose uptake for each region (see *Methods* for details). A negative LI indicated lower ipsilateral metabolism than contralateral, whereas a positive LI indicated higher ipsilateral metabolism than contralateral. This relative definition of hypo-vs. hypermetabolism is commonly used in clinical settings^46^. Leveraging this measure of metabolic integrity, we sought to identify the neurophysiological basis of the energetic inefficiency observed in our TLE patients.

Our first and most straightforward observation was that all the regions with disrupted energetic profiles captured by our NCT analyses — ipsilateral temporal pole, inferior temporal gyrus, amygdala, hippocampus, parahippocampal gyrus, fusiform gyrus, and isthmus of cingulate gyrus — also exhibited ipsilateral hypometabolism (permutation-based one-sample *t*-test correcting for multiple comparisons, **Supplementary Table 1**, **Figure 5a**). To expound on this observation, we also calculated LIs of the regional MCE in these regions, and tested bivariate correlations between the 7 pairs of LIs (i.e., one for MCE and one for glucose uptake) with a permutation-based product-moment correlation controlling for multiple comparisons. We found significant correlations between LIs at the temporal pole (*R*_49_=-0.37, *P*_corr_=0.049), amygdala (*R*_49_=-0.62, *P*_corr_=8×10^−6^), hippocampus (*R*_49_=-0.60, *P*_corr_=3×10^−5^), parahippocampal gyrus (*R*_49_=-0.50, *P*_corr_=0.002), and the fusiform gyrus (*R*_49_=-0.39, *P*_corr_=0.036), but not within the inferior temporal gyrus (*R*_49_=-0.34, *P*_corr_=0.096) or isthmus of cingulate gyrus (*R*_49_=-0.22, *P*_corr_=0.551) (**Figure 5b-h**). These results suggest that the regions where TLE patients show greater control energy needs also show greater *hypo*metabolism (*i*.*e*., lower metabolic baseline) with respect to their contralateral counterparts.

**Figure 5.**
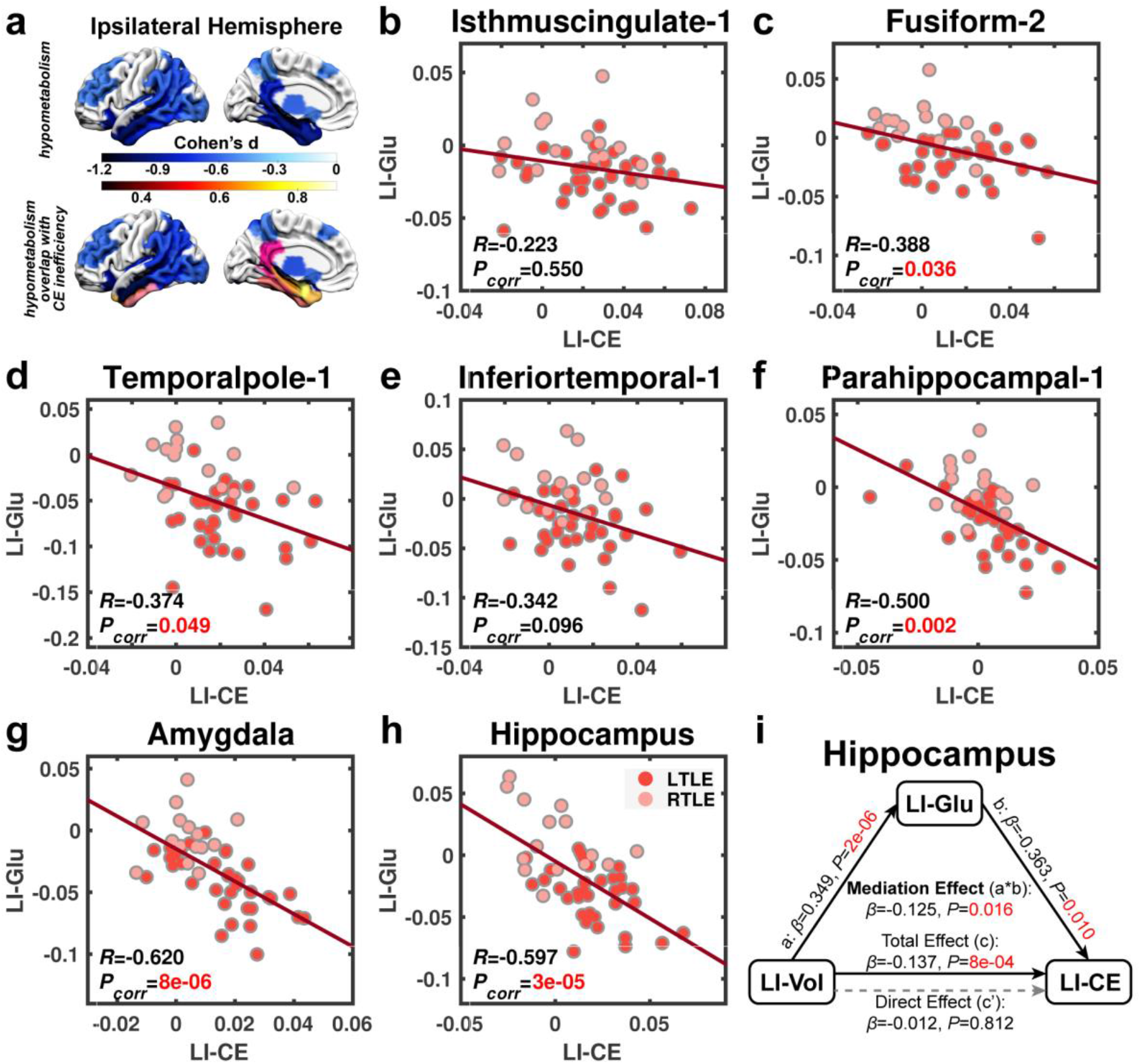
Regional control energy consumption is associated with glucose metabolism in temporal lobe epilepsy patients. (a) Multiple comparisons corrected one sample *t*-tests on laterality indices (LI) of regional glucose uptake reveal widespread ipsilateral hypometabolism in reference to the metabolic levels in the contralateral hemisphere (upper panel). Notably, all the ipsilateral temporo-limbic regions with atypical energetic profiles also present with hypometabolism (lower panel). (b-h) Pearson correlations corrected for multiple comparisons revealed significant associations between the laterality of glucose uptake and minimal control energy consumption during all the transitions between the states defined by intrinsic connectivity networks (ICN), most prominently in the limbic regions, whereas the side with lower glucose metabolic baseline consumes more control energy. Corrected *P*-values (*P*_corr_) are depicted. (i) A mediation analysis is performed on the hippocampus, where the association between the laterality of gray matter volume (LI-Vol) and control energy (LI-CE) is also found to be significant. We found that the laterality of glucose uptake (LI-Glu) provides a full mediation of the association between the former two variables. The significance of the mediation effect was assessed using bootstrapped confidence intervals.

One common reason for a region to exhibit hypometabolism is the loss of local structural integrity as manifest by gray matter atrophy^47,48^. Accordingly, within the above-mentioned regions, we also examined whether the LI of control energy was correlated with the LI of gray matter volume. We found a significant correlation between energy and volume LIs in the hippocampus only (*R*_49_=-0.47, *P*_corr_=0.005; all other regions had |*R*_49_|’s<0.32 and *P*_corr_’s>0.167). This finding indicates that greater gray matter volume loss in the hippocampus is associated with greater control energy needs. Interestingly, a bootstrapped mediation analysis focusing on the hippocampus found that the LI of glucose uptake provided full mediation of the association between LI of gray matter volume and LI of control energy (*β*=-0.125, *P*=0.016, **Figure 5i**). Thus, the loss of local structural integrity may serve as a substrate leading to a decline in baseline regional metabolism, which in turn engenders inefficient control of brain dynamics.

## Discussion

Pathological disruptions to normal brain dynamics in patients with drug-resistant epilepsy are associated with both transient and enduring detrimental effects on brain function^10,49^, which can significantly compromise quality of life. Developing treatments to alleviate such impairments is challenging owing to our limited understanding of how pathological conditions such as TLE alter energetic processes needed to facilitate desired brain dynamics. To this end, the recent development of network control theory has provided new opportunities to profile the energy landscape of the brain during simulated neural dynamics in which activity spreads along the structural connectome. Leveraging this framework, we showed that TLE patients required more control energy to activate the limbic network compared to HCs. In particular, this increased energy was localized to the limbic network ipsilateral to patients’ seizure focus. Further, we quantified regional energy profiles during transitions between different brain states, and found that the mesial and inferior parts of the ipsilateral temporal lobe in TLE consumed more control energy on average. Intuitively, the extra energetic demands in these patients may result in suboptimal dynamics and inadequate activation, and eventually impair function. Additionally, by conceptualizing TLE as a lateralized lesion model, we demonstrated that the imbalance of energetic costs between the ipsilateral and contralateral mesial and inferior temporal regions was also mirrored in their asymmetric metabolic patterns, whereby regions with lower baseline metabolic levels also had higher energetic demands. Specifically, for the hippocampus, we found significant associations between lateralization of energy costs, glucose uptake and gray matter volume, with hypometabolism fully mediating the increase in energy demand pertaining to volume loss on the ipsilateral side.

In this study, we focused on two main categories of brain state transitions. The first probed the efficiency with which patients’ brains could attain each of 8 ICN-defined preferential brain states from a common baseline. Compared to HCs, TLE patients needed greater global control energy to activate the limbic network, especially its ipsilateral side. Thus, it is plausible that failing to meet these increased energy demands may underpin inadequate limbic activation and dysfunction. Considering the overlap between the limbic network and the seizure focus in TLE, dysfunction within this network is expected in these patients. For instance, episodic memory deficits and affective comorbidities are commonly reported in TLE^11,13^, and can be reasonably attributed to limbic dysfunction. Previous imaging studies have shown that some limbic regions, such as the epileptogenic hippocampus, are less activated during episodic memory encoding in TLE^50–52^. Finally, altered functional connectivity seeded from the amygdala is also associated with comorbid psychiatric symptoms in these patients^53^. In line with this evidence, our findings suggest that TLE is associated with compromised energetic efficiency of ipsilateral limbic regions.

Brain function depends on the ability to reach desired brain states and to swiftly transition among them. Therefore, our second set of analyses focused on modeling the brain’s capacity to transition between pairs of ICN-defined preferential states. Across all possible pairwise transitions, we found that TLE patients exhibited elevated control energy requirements compared to HCs, again rooted in ipsilateral temporo-limbic regions. These results suggest that disruption to the underlying structural networks of TLE patients not only affects the activation of the limbic network, but also leads to greater energy demands of ipsilateral temporo-limbic regions during transitions among all ICN-defined states. Finally, we sought to further validate our findings by testing 100,000 pairs of random brain states that were not *a priori* rooted in functional neuroanatomy. In doing so, we found near-identical results, demonstrating that the increased energy costs in the ipsilateral temporo-limbic regions represent a general signature of dysfunctional control of neuronal dynamics in TLE patients. Collectively, our study provides evidence that altered brain dynamics in TLE, as a pathological trait, are underpinned by energetic inefficiency that mostly affects areas in proximity or closely connected to the seizure focus, and may represent a substrate of enduring brain dysfunction.

What is the neurophysiological basis of this trait? While NCT has been applied to neuroscience in recent years, the biological nature of control energy has remained unclear. We know that the brain consumes energy via glucose metabolism^6^, and previous studies have linked control energy to cognitive effort^25^. Therefore, we hypothesized that the magnitude of control energy may reflect the extent of local metabolism needed to instantiate the desired neural dynamics. Indeed, using FDG-PET, we showed that regional energetic inefficiency coexists with altered metabolism in TLE. Considering unilateral TLE as a lesion model, we observed that reduced baseline glucose intake (*i*.*e*., hypometabolism) aligned, as clinically expected, with the side of seizure focus. Taking the contralateral hemisphere as reference, we found that greater ipsilateral hypometabolism was associated with greater ipsilateral energetic inefficiency. For the first time, we thus highlight a formal link between theoretical control energy and a physiological measure of brain metabolism, and suggest that the compromised metabolic baseline in affected regions may lead to greater energetic challenges in supporting desired brain dynamics.

A common cause of metabolic alterations may be the loss of underlying structural integrity. For instance, concomitant gray matter volume loss and hypometabolism is reported in patients with TLE, especially in epileptogenic regions such as the hippocampus^54^. In our TLE patients, we found that gray matter volume asymmetry was also associated with energy asymmetry in the hippocampus. Through a mediation analysis, we formally demonstrated that the asymmetry of baseline metabolism fully mediates the association between the asymmetry of gray matter volume and energy costs. That is, greater volume loss may lead to greater baseline hypometabolism, thereby increasing energy demands during brain state transitions in the ipsilateral hippocampus. Such results deliver a unifying framework, linking independently measured regional volumetrics, glucose metabolism, and network control properties derived from structural networks. Our work thus captures both the metabolic and volumetric bases of control energy, further supporting its application in modeling the endogenous resources consumed during brain dynamics in the absence of external stimulation. In addition, our work suggests that the magnitude of control energy is not only modulated by the transition trajectory^25,29,37^, but also by the integrity of the underlying structure. Specifically, regions harboring pathology, such as the hippocampus in TLE, can exhibit different degrees of neural loss, causing a metabolic resource gap that in turn impacts brain state transitions.

Several methodological considerations are pertinent to this study. First, the structural connectome obtained via diffusion tractography used in our study is an approximation of the real structural scaffold of functional brain dynamics. Implementing other forms of structural connectivity, such as a spatial adjacency network, may provide added value to our model^37^. Second, we modeled the neural dynamics under assumptions of linearity and time-invariance, following previous studies^26,29,36,37,40^. Recent research has shown that such simplified models can provide useful first-order approximations of brain dynamics^55,56^, and even outperform non-linear models when predicting the macroscopic brain activity measured by functional magnetic resonance imaging and intracranial electrocorticography^57^. Nonetheless, further adaptations are expected to incorporate advanced features such as non-linearity^58,59^ and time-dependence^60^. Third, as in previous studies^26,36,40^, we defined a discrete set of brain states based on ICNs known to underpin cognition^32,34,35^. Alternatively, the magnitude of brain states could also be defined via empirically measured neurophysiological signals^25,27,29^. However, the estimated energy costs in our model not only depend on the network structure, but also on the distance between the initial and final states^29^. Thus, by setting binary states uniformly, we ensured a consistent transition distance across all subjects. Accordingly, the extent of energy costs only reflects the efficiency of the underlying network structure (or the lack thereof) during the same designated dynamic process. Fourth, our TLE cohort was heterogeneous in etiology, which may raise the possibility of subgroup-specific energetic characteristics that were not addressed in the current study. Last, some antiepileptic drugs (AED) may affect brain dynamics^61^; however, due to the heterogeneous regimen of AED history in our patients, we did not focus on the relationship between AED and control energy profiles here. Similarly, while interictal epileptic discharges (IEDs) can transiently influence brain dynamics, their relevance to control energy warrants further investigation, as quantitative measures of IEDs were not available in this study.

## Outlook

Focusing on temporal lobe epilepsy as a disease model, our study provides a framework linking loss of structural integrity, alteration of local metabolism, and greater energetic challenges to attain desired brain state transitions, leading to altered brain dynamics. By providing a neurophysiological basis of control energy, our work paves the way for further applications of network control theory in the field of neuroscience.

## Methods

### Participants

Sixty patients with refractory unilateral TLE (38 left-sided, 22 right-sided) were recruited from the Thomas Jefferson Comprehensive Epilepsy Center. Diagnosis was determined by a multimodal evaluation including neurological history and examination, interictal and ictal scalp video-EEG, MRI, FDG-PET, and neuropsychological testing. Localization was determined after confirming that the testing was concordant for unilateral temporal lobe epilepsy, as described previously^62^. Patients were excluded from the study for any of the following reasons: previous brain surgery, evidence for extra-temporal or multifocal epilepsy by history or testing, contraindications to MRI, or hospitalization for any Axis I disorder listed in the DSM-5 (Diagnostic and Statistical Manual of Mental Disorders, V). Depressive disorders were admissible, given the high comorbidity of depression and epilepsy^63^. The demographic and clinical characteristics of the patient groups are presented in ***Table 1***, along with the demographic information of 50 age-, sex-, handedness-, and education-matched healthy controls (HCs). All HC were free of psychiatric or neurological disorders based on a health screening measure. This study was approved by the Institutional Review Board for Research with Human Subjects at Thomas Jefferson University. All participants provided informed consent in writing.

### Imaging Acquisition

All participants were scanned on a 3-T X-series Philips Achieva clinical MRI scanner (Amsterdam, the Netherlands) at Thomas Jefferson University Hospital. The data acquisition session included both a High Angular Resolution Diffusion Imaging (HARDI) scan as well as a high-resolution T1-weighted (T1w) anatomical scan. The HARDI scan was 61-directional with a b-value of 3000 s/mm^2^ and TE/TR = 7517/98 ms, in addition to 1 b0 images. Matrix size was 96 × 96 with a slice number of 52. Field of view was 230 × 230 mm and slice thickness was 2.5 mm. Participants lay in a foam pad to comfortably stabilize the head, and were instructed to remain still throughout the scan. Prior to collection of the HARDI scan, T1w images (180 slices) were collected using an MPRage sequence (256 × 256 isotropic 1mm voxels; TR = 640 ms, TE = 3.2 ms, flip angle = 8°, FOV = 256 × 256 mm) in identical positions to provide an anatomical reference. The in-plane resolution for each T1 slice was 1 mm^3^ (axial oblique).

As part of their presurgical evaluation, 50 patients also underwent on-site PET scans. The other 10 patients who received PET scans from other facilities (off-site), and HC who did not receive PET scans, were excluded from the FDG-PET related analysis. All PET scans were performed during interictal periods using a standard protocol. Pre-injection blood glucose level was below 150 mg/dl for all patients (range: 61–128 mg/dl). An intravenous catheter was inserted under local anesthesia and a dose around 5.9 ± 1.4 mCi of radioactive 100mg/l fluorodeoxyglucose (FDG) was injected. The scan was initiated about 42 ± 15 min after the injection. Participants’ eyes were open, and their ears were non-occluded. Ambient noise and light was kept to a minimum. Thirty-one patients (62% of patients; 20 LTLE, 11 RTLE) were scanned on a Siemens Biograph 1080 PET/CT, with data consisting of 109 axial slices, 3 mm thick, and 1 × 1 mm in resolution. The remaining 19 patients (38% of patients; 15 LTLE, 4 RTLE) were scanned on a Siemens Biograph 20 mCT PET/CT, with data consisting of 110 axial slices, 3 mm thick, and 1.6 × 1.6 mm in resolution. Attenuation-corrected PET images were iteratively reconstructed by standard vendor-provided software. There was no significant difference in proportion of left and right TLE patients acquired with either scanner (*χ*^2^ = 1.17, p = 0.28). Furthermore, we obtained asymmetry indices that use the same subject as reference, therefore reducing potential scanner-specific bias, as reported previously^64^. This procedure also avoids confounds related to demographic factors, such as age, medication history, and epilepsy history^65,66^. Nevertheless, the scanner model was also used as a categorical nuisance regressor during the data analysis along with demographic information.

### Imaging Processing

The T1w and HARDI data were analyzed through QSIprep^67^ (v0.8.0, https://qsiprep.readthedocs.io), which is based on Nipype 1.4.2. QSIprep automates diffusion MRI data preprocessing and reconstruction using well-recognized neuroimaging tools including Advanced Normalization Tools (ANTs), Analysis of Functional NeuroImages (AFNI), FMRIB Software Library (FSL), DSI Studio^68^, MRtrix 3^69^, and fMRIprep^70^.

#### Anatomical data preprocessing

The T1w image was corrected for intensity non-uniformity using N4BiasFieldCorrection (ANTs 2.3.1^71^), and was used as a T1w-reference throughout the workflow. The T1w-reference was then skull-stripped using antsBrainExtraction.sh (ANTs 2.3.1), using OASIS as the target template. Spatial normalization to the ICBM 152 Nonlinear Asymmetrical template version 2009c was performed through nonlinear registration with antsRegistration (ANTs 2.3.1^72^), using brain-extracted versions of both T1w volume and template. Brain tissue segmentation of cerebrospinal fluid (CSF), white-matter (WM) and gray-matter (GM) was performed on the brain-extracted T1w using FAST (FSL 6.0.3^73^).

#### Diffusion data preprocessing

The HARDI image was first denoised using a Marchenko-Pastur principal component analysis (MP-PCA) method^74^, then underwent Gibbs-ringing artifacts removal^75^, and then was spatially bias corrected through MRtrix 3. Subsequently, it was corrected for head motion and eddy current distortions via eddy_openmp (FSL 6.0.3). A deformation field to correct for susceptibility distortions was estimated based on fMRIprep’s fieldmap-less approach. The deformation field was that resulting from co-registering the b0 reference to the same-subject T1w-reference with its intensity inverted^76^. Registration was performed with antsRegistration (ANTs 2.3.1), and the process was regularized by constraining the deformation to be nonzero only along the phase-encoding direction, and was further modulated with an average fieldmap template^77^. Based on the estimated susceptibility distortion, an unwarped b=0 reference was calculated for a more accurate co-registration with the T1w-reference, and then a final preprocessed HARDI image was produced in the native space of the T1w-reference with an isotropic voxel size of 2 mm^3^. Two quality metrics were calculated based on the preprocessed data, including framewise displacement^78^ and neighboring correlation^79^.

#### Diffusion data reconstruction and tractography

The preprocessed HARDI image was reconstructed via MRtrix 3. Multi-tissue fiber response functions were estimated using the Dhollander algorithm, during which Fibre Orientation Distributions (FODs) were estimated via constrained spherical deconvolution [CSD^80,81^] using an unsupervised multi-tissue method^82,83^. Specifically, a single-shell-optimized multi-tissue CSD was performed using MRtrix3Tissue (https://3Tissue.github.io), a fork of MRtrix3. FODs were intensity-normalized using mtnormalize^84^. Subsequently, an anatomically-constrained probabilistic tractography^85^ was performed using the iFOD2 probabilistic tracking method, in which the WM FODs were used for tractography and the T1w segmentation was used for anatomical constraints. For each subject we generated 10^7^ streamlines with a maximum length of 250 mm, minimum length of 30 mm, and FOD power of 0.33. Weights for each streamline were calculated using a spherical-deconvolution informed filtering of tractograms (SIFT2)^86^, which was then used to estimate the structural connectivity matrix.

#### Brain parcellation customization

Consistent with previous work^36^, we chose the Lausanne parcellation scheme including n = 129 cortical and subcortical parcels^87^ to build the structural network. This parcellation scheme is established by sub-dividing the Desikan-Killiany anatomical atlas^88^. Specifically, to enable the proposed asymmetry analysis, we needed the parcellation to be symmetric. However, we noted that 6 regions have been subdivided asymmetrically, whereas the medial orbito-frontal gyrus, inferior parietal gyrus, and lateral occipital gyrus have one more subdivision in the right hemisphere, and the rostral middle frontal gyrus, precentral gyrus, postcentral gyrus have one more subdivision in the left hemisphere. These additional subdivisions were subsequently merged with their corresponding neighbor to match their cross-hemisphere counterpart, producing a symmetric version of the parcellation constituted by 61 pairs of cortical and subcortical parcels (excluding brainstem; details in ***Supplementary Table 2***). This parcellation was then inversely warped onto the native space of the T1w-reference and resampled at 2 mm^3^ voxel resolution. Using tck2connectome (MRTrix3) allowing a 2 mm radial search from each streamline endpoint to locate the nearest node, we built a 122 × 122 undirected adjacency matrix for each subject with the SIFT2 weighted streamline counts representing interregional structural connectivity.

To define neurobiologically meaningful brain states, we capitalized on an established functional brain parcellation^89^ of intrinsic connectivity networks (ICNs), which was defined by clustering the resting-state functional connectivity data from 1000 healthy subjects^33^. This parcellation is constituted by 7 cortical ICNs that are commonly seen during both resting and task conditions^32,34^, including visual (VIS), somatomotor (SM), dorsal attention (DAN), salience/ventral attention (SAL/VAN), limbic (LIM), executive control (CONT), and default mode networks (DMN). As in prior work^36^, we mapped both parcellations to a common surface space (fsaverage), and calculated the proportional overlap of vertices between each parcel and each of the 7 ICNs. Using a winner-take-all strategy, we assigned each parcel to the ICN with highest overlap proportion (**Figure 1b**). In addition, subcortical regions were summarized in an eighth, subcortical network (SC), excepting for the hippocampus and amygdala, which were assigned to the limbic network following the common clinical definition^14^. These 8 non-overlapping networks were used as representative brain states during simulations of brain state transitions.

#### FDG-PET data preprocessing

PET images were preprocessed with Statistical Parametric Mapping 12 (SPM 12, http://www.fil.ion.ucl.ac.uk/spm/software/spm12), as in previous work^90^. Briefly, the PET image was first co-registered to the T1w-reference image, smoothed with a 6-mm kernel, and intensity-normalized by the global mean uptake estimated based on a skull-striped brain mask derived from the T1w-reference image. Regional mean glucose uptake was subsequently estimated based on the same parcellation. As stated, in the absence of PET data from HC, we calculated the laterality index 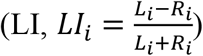 of regional glucose uptake from the aforementioned 61 pairs of parcels for the subsequent analyses.

#### Regional gray matter volume estimation

Lastly, we obtained regional mean gray matter volumes (GMV) using the Computational Anatomy Toolbox (CAT12, v12.7, http://www.neuro.uni-jena.de/cat/). The T1w image was first denoised with a spatial adaptive non-local means (SANLM) denoising filter^91^, followed by internal resampling to properly accommodate low-resolution images and anisotropic spatial resolutions. Subsequently, the data was bias-corrected and affine-registered followed by the standard SPM “unified segmentation”^92^. The brain was then parcellated into left and right hemispheres, subcortical areas, and the cerebellum. Furthermore, local white matter hyperintensities were detected to be later accounted for during the spatial normalization. Subsequently, a local intensity transformation of all tissue classes was performed, and a final adaptive maximum *a posteriori* (AMAP) segmentation^93^ was then refined by applying a partial volume estimation^94^, which effectively estimated the fractional content for each tissue type per voxel. Last, the tissue segments were spatially normalized to a common reference space using Geodesic Shooting^95^ registration, so that the regional GMV could be estimated based on the same parcellation. In addition, the total intracranial volume and a summary image quality rating for each T1w image were exported, and were used as covariates.

### Brain State Transitions Simulated through Linear Network Control Theory

#### Theoretical framework of Linear Network Control Theory

To investigate whether TLE is associated with compromised efficiency of common brain dynamics, we leveraged recent developments of linear network control theory, and explored the energetic efficiency of the structural brain network in facilitating designated brain state transitions. As in previous work^26,29,36,37,40^, we employed a simplified noise-free linear and time-invariant network model to describe the dynamics of the brain:

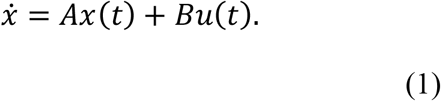

Here, *x*(*t*) is a N × 1 vector that represents the state (*i*.*e*., activity level) of each node of the system at time *t* (N = 122). *A* is a N × N adjacency matrix denoting the relationship between the system elements, which can be operationalized as the structural brain network. To ensure the stability of the system, *A* is normalized as follows^37^:

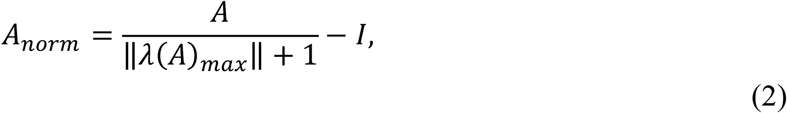

whereas *λ*(*A*)_*max*_ denotes the largest positive eigenvalue of the system and *I* denotes the N × N identity matrix. *B* is the input matrix that identifies the nodes in the control set. Here we set *B* to be the N × N identity matrix to set all the brain parcels as control nodes where energy can be consumed to facilitate brain state transitions. Last, *u*(*t*) denotes the amount of energy injected into each control node at each time point *t*. Intuitively, *u*(*t*) can be summarized over time to represent the total energy consumption during transition from an initial state to a final state.

#### Simulation I: Individual ICN activation

In our first simulation, we considered the scenario that the brain transits from an initial baseline state (*x*_*0*_ = *x*(*t* = 0)) to a final state (*x*_*T*_ = *x*(*t* = *T*)) when a specific ICN is predominantly activated. We modeled this control task by setting:

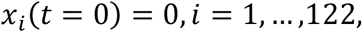

and

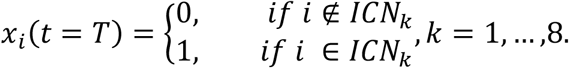

Note that, theoretically, setting the initial state to full zeros does not necessarily mean that the brain is globally inactive, which is biologically impossible. Rather, it can be viewed as a mean-centered baseline, and the final state has additional activations within the specific ICN than other regions by 1 arbitrary unit. This setting is analogous to task fMRI analyses, where contrasts are commonly set between a condition of interest (1) and a baseline (0)^26^.

To explore the energetic efficiency of the structural brain network in facilitating the activation of specific ICNs, we adopted the optimal control framework to estimate the control energy required to optimally steer the brain through these state transitions^26,29^. Against a naturalistic trajectory, when the brain state drifts without any control input, the proposed state transition commonly relies on the additional control input *u*(*t*) to reach the desired final state. This control effort can be intuitively understood as an internal cognitive control process (or as external brain stimulation in other patient scenarios), and it is based on both the energy costs and the length of the transition trajectory^37^. Therefore, an optimal solution can be described as the minimized combination of both the length of the transition trajectory and the required control energy, during the state transition from an initial state (*x*(0) = *x*_0_) to the final state (*x*(*T*) = *x*_*T*_) over the time horizon *T* (see Refs. ^40,96^):

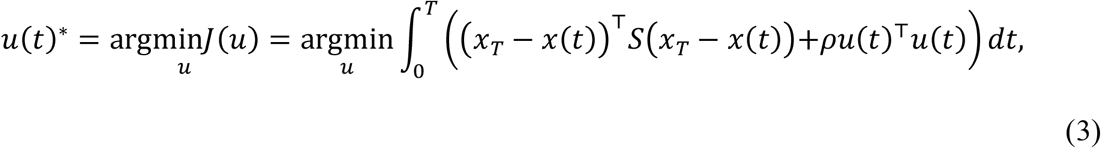

where (*x*_*T*_ − *x*(*t*)) ^T^(*x*_*T*_ − *x*(*t*)) is the distance between the state at time *t* and the final state *x*_*T*_, *T* is the finite amount of time given to reach the final state, and *ρ* is the relative weighting between the cost associated with the length of the transition trajectory and the input control energy. Following a previous benchmarking study^37^, we set *T =* 3 and *ρ =* 1, allowing 1000 steps of transition from the initial state to the final state. To minimize the unintended energy cost on regulating the regions of no interest (*i*.*e*., those outside of the target ICN), we applied a constraint matrix *S*, which is a N × N binary diagonal matrix that selects only regions that are members of the target ICN. Accordingly, the term (*x*_*T*_ − *x*(*t*)) ^T^ *S*(*x*_*T*_ − *x*(*t*)) specifically constrains the trajectories of all regions within the target ICN, and the term *u*(*t*)^T^*u*(*t*) constrains the amount of control energy used to reach the final state. The cost function *J*(*u*(*t*)^*\**^) is used to solve the unique optimal control input *u*(*t*)^*\**^. Specifically, we define a Hamiltonian as:

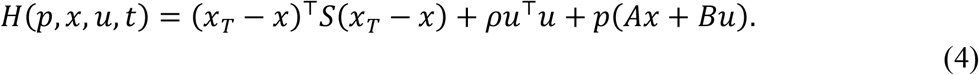

According to the Pontryagin minimization principle^96^, if *u*^*\**^ is a solution with the optimal trajectory *x*^*\**^, then there exists a *p*^*\**^ such that:

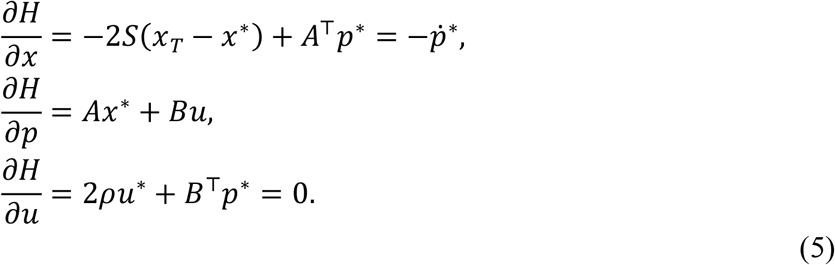

From this set of equations, we can derive that:

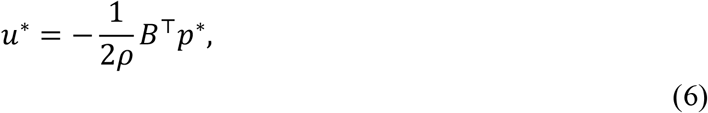

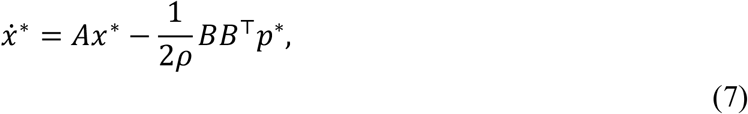

which can be reduced to:

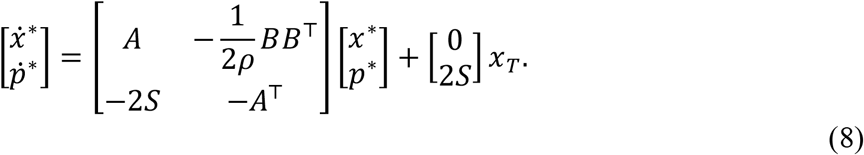

If we denote:

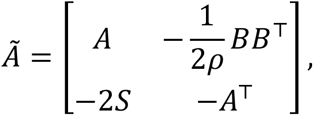

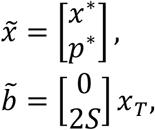

then this reduced equation (8) can be rewritten as:

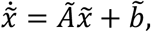

and can be solved as:

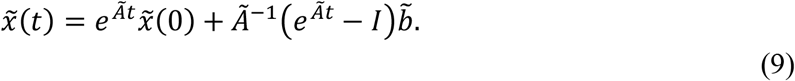

Then, by fixing *t = T*, we arrive at:

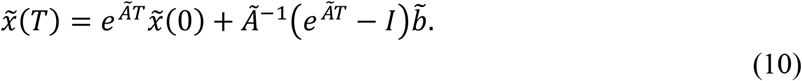

We let:

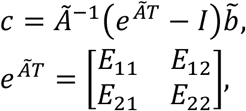

so that equation (10) can be rewritten as:

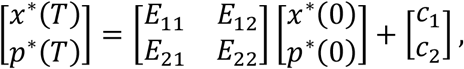

from which we can obtain:

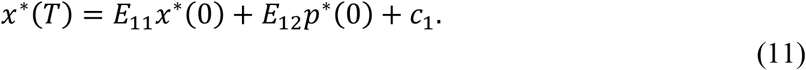

Moreover, if we let 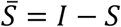, then as a known result in optimal control theory^24^, 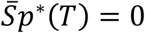. Therefore,

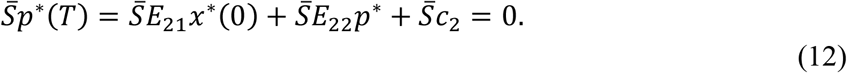

Finally, *p*^*\**^(*0*) can be solved for as follows:

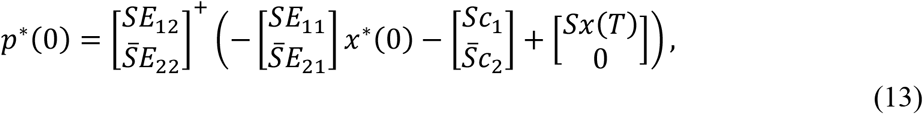

where [*·*]^*+*^ indicates the Moore-Penrose pseudoinverse of a matrix. Now that we have obtained *p*^*\**^(*0*), we can use it and *x(0*) to solve for 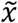 via forward integration based on equation (9). To solve for *u*^*\**^, we take *p*^*\**^ from our solution for 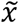 and solve equation (6). In particular, the optimal control energy injected at each region *i* can be defined as:

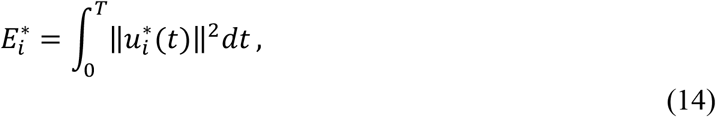

or given in total:

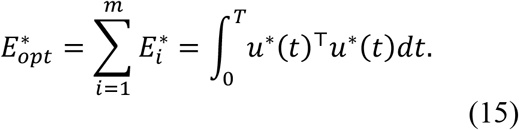

This total optimal control energy consumption 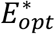 is then used as a measure of efficiency of the structural brain network during specific ICN activations.

#### Simulation IIa: Between ICN transitions

In our second simulation, we investigated the regional energetic consumption associated with facilitating brain dynamics. While it is computationally impossible to simulate all brain transitions, we considered two sets of finite repositories instead. First, we used the 8 ICN-defined representative brain states (*i*.*e*., the *x*_*T*_ in *Simulation I*), and explored all the possible transitions among them^36^. Counting scenarios of both reciprocal transitions and single state persistence (*i*.*e*., *x*_*0*_ *= x*_*T*_), this simulation resulted in a total of 64 control tasks. Considering the linear nature of our dynamical model, theoretically any possible transition can be written as a linear combination of the proposed transitions^36^. Thus, this simulation is generally relevant to all transitions represented during common brain dynamics.

We further alleviated the constraint on the length of the transition trajectory in our model, to allow the brain to travel more freely across different intermediate states. Specifically, we adopted a subform of the optimal control framework, namely the minimal control energy, which can be obtained by letting *ρ* → ∞ in equation (3), so that the cost function *J* accounts only for the control input to facilitate the designated transition regardless of the trajectory^37^. Accordingly, the minimal control energy during the state transition from an initial state (*x*(0) = *x*_0_) to the final state (*x*(*T*) = *x*_*T*_) over the time horizon *T* can be described as^4,27,37^:

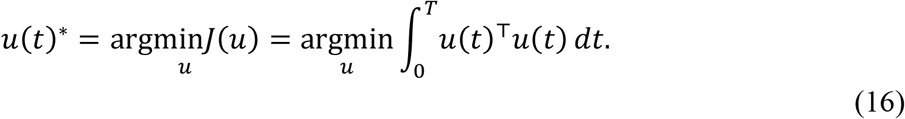

To solve the minimal control energy *u*(*t*)^*\**^ this time, we compute the controllability Gramian *W* for controlling the network *A* from the control node set *B* in equation (1) as:

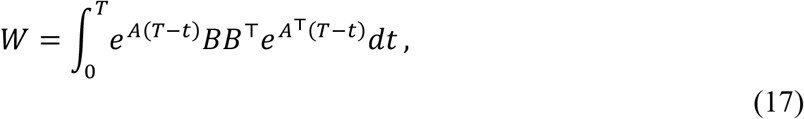

where, as defined previously, *A* is the normalized N × N structural brain network, *B* is a N × N identity matrix setting all the brain parcels as control nodes, and *T* is the finite time horizon of the transition trajectory. Similarly, we set *T =* 3 and allow for 1000 steps of transition from the initial state to the final state following Ref.^37^. Then, the *u*(*t*)^*\**^ can be computed as:

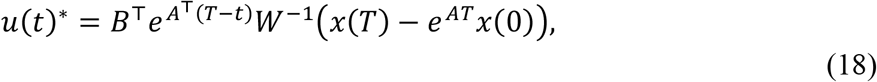

and the minimal control energy injected at each region *i* can calculated based on equation (14). Finally, for each brain region, we averaged their minimal control energy across the 64 control tasks as a measure of their individual energetic consumption during dynamics among known ICNs.

#### Simulation IIb: Random brain state transitions

The second set of finite repositories of brain states included 100,000 pairs of randomly generated initial and final brain states *x*_*rand*_ with a Gaussian distribution at *mean*(*x*_*rand*_) = 1 and *std*(*x*_*rand*_) = 0.1^26^. Accordingly, this simulation resulted in a total of 100,000 control tasks, which served as an approximation of all transitions when no prior preference of brain states is explicitly defined. Based on our previous argument of the linear nature of the model, we were not expecting significant difference between our previous *Simulation IIa* and this *Simulation IIb*. Rather, we expected to observe a consistency between the two, which would serve as a validation of *Simulation IIa*. Similarly, we adopted the same minimal control framework, and for each brain region, we calculated and summarized their minimal control energy across the 100,000 control tasks as the measure of their individual energetic consumption during brain dynamics among randomly organized brain networks.

### Statistical Inferences

Comparisons for common demographic and clinical information were made with standard parametric tests such as an independent *t*-test or Chi-Square test, conducted using IBM® SPSS® v25. The alpha level was set at *P* < 0.05 for both parametric and nonparametric tests.

#### Confounding factor regression

To minimize the influence of individual variances of the demographic characteristics and imaging data qualities (**Table 1**), confounding factor regressions were applied before statistical inferences on our neuroimaging data. Specifically, for derivates from all imaging modalities, 4 potential confounding factors were identified and included in the models: age, sex, handedness, and total intracranial volume. Furthermore, additional confounding factors were added to the models for different modalities: (i) for HARDI derivates (*i*.*e*., control energy), the neighboring correlation, mean framewise displacement, matrix density, and total weight were added; (ii) for T1w derivates (*i*.*e*., regional gray matter volume), the image quality rating was added; (iii) for FDG-PET derivates (*i*.*e*., regional glucose uptake), the PET scanner model was added. For each modality, all confounding factors were regressed out from their derivates with one linear regression model, and the residuals were taken for subsequent statistical analyses.

#### Permutation-based Nonparametric Statistical Testing

To minimize the bias of the data distribution to our statistical inference and to correct for multiple comparisons, we implemented a permutation-based method as our main statistical strategy^41^. Individual permutation-based statistical testing allows inference of the probability of the observed statistic (*e*.*g*., *t*-value), from a distribution of the same statistic estimated from massive instances of the same samples with their group identities permuted^97^. In many cases, we wish to apply a permutation-based test to scenarios with multiple comparisons, *i*.*e*., comparing multiple within-subject variables across the same groups of subjects. In this case, we can expand the traditional approach by applying a “*t*_max_” principle to adjust the estimated *P*-values of each variable for multiple comparisons by controlling the family-wise error rate^42^. Briefly, the observed statistic for each variable is compared to the distribution of the most extreme statistic across the entire family of tests for each possible permutation. This procedure corrects for multiple comparisons, because the distribution of the most extreme statistics automatically adjusts to reflect the increased chance of false discoveries due to an increased number of comparisons^41^. We performed 1,000,000 permutations each time to ensure high precision during *P*-value estimation^44^. This strategy was applied on all analyses (*i*.*e*., *t*-tests, product-moment correlation) when multiple comparison correction was required.

To increase statistical power^38,43,44^, the regional control energy values of the right TLE patients were flipped left to right, allowing all statistical analyses to be conducted in accordance with the site of ictal onset (left, ipsilateral; right, contralateral). However, as there was no way to flip HC data to match the ipsilateral versus contralateral side in the right TLE patients, we instead calculated the deviation score of regional energy [*Z*_*pat*_ = (*E*_*pat*_ − *μ*_*con*_)/σ_*con*_, where *μ*_*con*_ and σ_*con*_ were the mean and standard deviation of the same regional energy from the HC] for each patient at each hemisphere, and flipped the Z-score of right TLE afterwards^38,44,45^. Then, Z-scores were evaluated via a permutation-based one-sample *t*-test.

#### Permutation-based Mediation Analysis

To disentangle the associations among regional gray matter volume change, glucose metabolism and control energy consumption in the hippocampus, we applied a mediation analysis to test the hypothesis that the regional metabolic baseline, as a measure of functional integrity, mediates the relationship between local structural integrity and energetic efficiency. After confound regression, we generated the laterality indices for gray matter volume, glucose uptake, and minimal control energy estimated during cross-ICN transitions. We then evaluated the significance of the indirect effect using bootstrapped confidence intervals via the MediationToolbox^98^. Specifically, we examined: (i) path c: the total effect of the LI of gray matter volume on the LI of minimal control energy; (ii) path a: the relationship between the LI of gray matter volume and the LI of glucose uptake; (iii) path b: the relationship between the LI of glucose uptake and the LI of minimal control energy; and (iv) path c’: the direct effect of the LI of gray matter volume on the LI of minimal control energy while controlling for the mediator (LI of glucose uptake). The mediation/indirect effect a*b is the effect size of the relationship between the LI of gray matter volume and the LI of minimal control energy that was reduced after controlling for the mediator (LI of glucose uptake). For each path, we calculated the beta coefficient, which reflected the changes of the outcome for every one-unit change in the predictor. A bootstrap analysis (*i*.*e*., resampled 1,000,000 times) was implemented to estimate the confidence intervals for the indirect effect.

## Data availability

The data are available from the authors upon reasonable request.

## Code availability

All codes are available from the authors upon reasonable request.

## Citation diversity statement

Recent work in several fields of science has identified a bias in citation practices such that papers from women and other minority scholars are under-cited relative to the number of such papers in the field^1-9^. Here we sought to proactively consider choosing references that reflect the diversity of the field in thought, form of contribution, gender, race, ethnicity, and other factors. First, we obtained the predicted gender of the first and last author of each reference by using databases that store the probability of a first name being carried by a woman^5, 10^. By this measure (and excluding self-citations to the first and last authors of our current paper), our references contain 7.41% woman(first)/woman(last), 18.52% man/woman, 18.52% woman/man, and 55.56% man/man. This method is limited in that a) names, pronouns, and social media profiles used to construct the databases may not, in every case, be indicative of gender identity and b) it cannot account for intersex, non-binary, or transgender people. Second, we obtained predicted racial/ethnic category of the first and last author of each reference by databases that store the probability of a first and last name being carried by an author of color^11, 12^. By this measure (and excluding self-citations), our references contain 9.79% author of color (first)/author of color(last), 12.4% white author/author of color, 25.2% author of color/white author, and 52.62% white author/white author. This method is limited in that a) names and Florida Voter Data to make the predictions may not be indicative of racial/ethnic identity, and b) it cannot account for Indigenous and mixed-race authors, or those who may face differential biases due to the ambiguous racialization or ethnicization of their names. We look forward to future work that could help us to better understand how to support equitable practices in science.

## Acknowledgements

We are grateful to Dr. Bianca De Blasi, Dr. Eli J. Cornblath, Dr. Matthew Cieslak, Dr. Pragya Srivastava, Dr. Richard F. Betzel, Dr. Urs Braun, and Dr. Zaixu Cui for useful discussions. D.S.B. acknowledges support from the Swartz Foundation, the Paul Allen Foundation, the Army Research Office (Bassett-W911NF-14-1-0679, Grafton-W911NF-16-1-0474), the National Institute of Mental Health (2-R01-DC-009209-11, R01-MH112847, R01-MH107235, R21-M MH-106799), National Institute of Neurological Disorders and Stroke (R01-NS099348), and the National Science Foundation (BCS-1631550 and IIS-1926757). X.H. acknowledges support from the Research Start-up Fund of USTC. LC acknowledges support from a Berkeley Fellowship, jointly awarded by University College London and Gonville and Caius College, Cambridge. LP was supported by the National Institute of Mental Health of the National Institutes of Health under Award Number K99MH127296 and a 2020 NARSAD Young Investigator Grant from the Brain & Behavior Research Foundation. J.I.T. acknowledges support from the National Institute of Mental Health (R01-MH104606) and National Institute of Neurological Disorders and Stroke (R01-NS112816-01). M.R.S. acknowledges support from the NIH and DARPA. J.Z.K. acknowledges support from the NSF Graduate Research Fellowship No. DGE-1321851. The content is solely the responsibility of the authors and does not necessarily represent the official views of any of the funding agencies.

## Author contributions

X.H. and D.S.B. designed the study. X.H. performed all analyses and wrote the initial draft of the manuscript. L.C. and L.P. contributed to the drafting and revising of the manuscript. J.S. and T.M.K. contributed to the analytical code. J.Z.K., Z.L., T.M., and F.P. contributed analytic solutions. M.R.S. and J.I.T. provided the clinical and imaging data. All authors edited the manuscript and approved the final version.

