## Supplementary Table for "Pathological and metabolic underpinnings of energetic inefficiency in temporal lobe epilepsy"

### Supplementary information

**Supplementary Table 1.** Permutation-based nonparametric one-sample *t*-tests on laterality indices (LIs) of regional glucose uptake reveal significant ipsilateral hypometabolism in temporal lobe epilepsy patients.

| Lausanne Atlas Region Label | LI (mean±std) | $t_{49}$ | $P_{corr}$ | Cohen's $d$ |
|---|---|---|---|---|
| lateralorbitofrontal_1 | -0.014±0.019 | -5.383 | $1\times10^{-4}$ | -0.761 |
| rostralmiddlefrontal_1 | -0.005±0.01 | -3.678 | 0.023 | -0.520 |
| superiorfrontal_3 | -0.007±0.013 | -3.622 | 0.027 | -0.512 |
| **isthmuscingulate_1** | -0.016±0.02 | -5.788 | $3\times10^{-5}$ | -0.819 |
| supramarginal_1 | -0.014±0.024 | -4.226 | 0.005 | -0.598 |
| superiorparietal_2 | -0.011±0.017 | -4.492 | 0.002 | -0.635 |
| inferiorparietal_1 | -0.013±0.019 | -4.690 | 0.001 | -0.663 |
| precuneus_2 | -0.017±0.036 | -3.401 | 0.049 | -0.481 |
| lateraloccipital_2 | -0.011±0.019 | -3.908 | 0.012 | -0.553 |
| **fusiform_2** | -0.016±0.019 | -6.196 | $1\times10^{-5}$ | -0.876 |
| **parahippocampal_1** | -0.018±0.021 | -5.981 | $2\times10^{-5}$ | -0.846 |
| entorhinal_1 | -0.039±0.042 | -6.652 | $5\times10^{-6}$ | -0.941 |
| **temporalpole_1** | -0.045±0.049 | -6.608 | $5\times10^{-6}$ | -0.935 |
| **inferiortemporal_1** | -0.024±0.03 | -5.695 | $4\times10^{-5}$ | -0.805 |
| inferiortemporal_2 | -0.013±0.017 | -5.452 | $1\times10^{-4}$ | -0.771 |
| middletemporal_1 | -0.012±0.02 | -4.191 | 0.005 | -0.593 |
| middletemporal_2 | -0.026±0.027 | -6.795 | $2\times10^{-6}$ | -0.961 |
| bankssts_1 | -0.02±0.023 | -6.245 | $1\times10^{-5}$ | -0.883 |
| superiortemporal_1 | -0.014±0.02 | -4.742 | 0.001 | -0.671 |
| insula_1 | -0.024±0.039 | -4.485 | 0.002 | -0.634 |
| thalamusproper | -0.008±0.013 | -4.228 | 0.005 | -0.598 |
| putamen | -0.005±0.008 | -4.830 | 0.001 | -0.683 |
| pallidum | -0.013±0.021 | -4.422 | 0.003 | -0.625 |
| accumbensarea | -0.008±0.015 | -3.746 | 0.019 | -0.530 |
| **hippocampus** | -0.026±0.026 | -7.173 | $1\times10^{-6}$ | -1.014 |
| **amygdala** | -0.028±0.031 | -6.423 | $9\times10^{-6}$ | -0.908 |

* The LI is calculated as $LI_i = \frac{Ipsilateral_i - Contralateral_i}{Ipsilateral_i + Contralateral_i}$, whereas a negative LI indicated lower ipsilateral compared to contralateral metabolism. Only regions with LIs significantly different from zero after multiple comparisons corrections are presented. Statistics including the *t*-value, corrected *P*-value ($P_{corr}$), and effect size (Cohen's *d*) are depicted. The regions co-exhibiting ipsilateral abnormalities in energy profiles are highlighted in bold.

**Supplementary Table 2.** Label correspondence between the symmetrically modified Lausanne Atlas and the original Lausanne Atlas[87]. Each region is assigned to one of the intrinsic connectivity networks (ICNs)[33] based on spatial overlap between the Lausanne Atlas and the Schaefer Atlas[89].

| **Original Lausanne Atlas** | | **Symmetrically modified Lausanne Atlas** | | **ICN** |
|---|---|---|---|---|
| *Index* | *Region Label* | *Index* | *Region Label* | |
| 1 | R_lateralorbitofrontal_1 | 1 | R_lateralorbitofrontal_1 | LIM |
| 2 | R_lateralorbitofrontal_2 | 2 | R_lateralorbitofrontal_2 | LIM |
| 3 | R_parsorbitalis_1 | 3 | R_parsorbitalis_1 | DMN |
| 4 | R_frontalpole_1 | 4 | R_frontalpole_1 | LIM |
| 5 | R_medialorbitofrontal_1 | 5 | R_medialorbitofrontal_1 | DMN |
| 6 | R_medialorbitofrontal_2 | 5 | R_medialorbitofrontal_2 | DMN |
| 7 | R_parstriangularis_1 | 6 | R_parstriangularis_1 | DMN |
| 8 | R_parsopercularis_1 | 7 | R_parsopercularis_1 | SAL/VAN |
| 9 | R_rostralmiddlefrontal_1 | 8 | R_rostralmiddlefrontal_1 | CONT |
| 10 | R_rostralmiddlefrontal_2 | 9 | R_rostralmiddlefrontal_2 | CONT |
| 11 | R_superiorfrontal_1 | 10 | R_superiorfrontal_1 | DMN |
| 12 | R_superiorfrontal_2 | 11 | R_superiorfrontal_2 | DMN |
| 13 | R_superiorfrontal_3 | 12 | R_superiorfrontal_3 | SAL/VAN |
| 14 | R_superiorfrontal_4 | 13 | R_superiorfrontal_4 | DAN |
| 15 | R_caudalmiddlefrontal_1 | 14 | R_caudalmiddlefrontal_1 | CONT |
| 16 | R_precentral_1 | 15 | R_precentral_1 | SAL/VAN |
| 17 | R_precentral_2 | 16 | R_precentral_2 | SMN |
| 18 | R_precentral_3 | 17 | R_precentral_3 | SMN |
| 19 | R_paracentral_1 | 18 | R_paracentral_1 | SMN |
| 20 | R_rostralanteriorcingulate_1 | 19 | R_rostralanteriorcingulate_1 | DMN |
| 21 | R_caudalanteriorcingulate_1 | 20 | R_caudalanteriorcingulate_1 | SAL/VAN |
| 22 | R_posteriorcingulate_1 | 21 | R_posteriorcingulate_1 | CONT |
| 23 | R_isthmuscingulate_1 | 22 | R_isthmuscingulate_1 | DMN |
| 24 | R_postcentral_1 | 23 | R_postcentral_1 | SMN |
| 25 | R_postcentral_2 | 24 | R_postcentral_2 | SMN |
| 26 | R_supramarginal_1 | 25 | R_supramarginal_1 | DAN |
| 27 | R_supramarginal_2 | 26 | R_supramarginal_2 | SAL/VAN |
| 28 | R_superiorparietal_1 | 27 | R_superiorparietal_1 | DAN |
| 29 | R_superiorparietal_2 | 28 | R_superiorparietal_2 | DAN |
| 30 | R_superiorparietal_3 | 29 | R_superiorparietal_3 | VIS |
| 31 | R_inferiorparietal_1 | 30 | R_inferiorparietal_1 | DMN |
| 32 | R_inferiorparietal_2 | 30 | R_inferiorparietal_2 | DMN |
| 33 | R_inferiorparietal_3 | 31 | R_inferiorparietal_3 | DAN |

| | | | | |
|---|---|---|---|---|
| 34 | R_precuneus_1 | 32 | R_precuneus_1 | DMN |
| 35 | R_precuneus_2 | 33 | R_precuneus_2 | DMN |
| 36 | R_cuneus_1 | 34 | R_cuneus_1 | VIS |
| 37 | R_pericalcarine_1 | 35 | R_pericalcarine_1 | VIS |
| 38 | R_lateraloccipital_1 | 36 | R_lateraloccipital_1 | VIS |
| 39 | R_lateraloccipital_2 | 37 | R_lateraloccipital_2 | VIS |
| 40 | R_lateraloccipital_3 | 37 | R_lateraloccipital_3 | VIS |
| 41 | R_lingual_1 | 38 | R_lingual_1 | VIS |
| 42 | R_lingual_2 | 39 | R_lingual_2 | VIS |
| 43 | R_fusiform_1 | 40 | R_fusiform_1 | VIS |
| 44 | R_fusiform_2 | 41 | R_fusiform_2 | LIM |
| 45 | R_parahippocampal_1 | 42 | R_parahippocampal_1 | LIM |
| 46 | R_entorhinal_1 | 43 | R_entorhinal_1 | LIM |
| 47 | R_temporalpole_1 | 44 | R_temporalpole_1 | LIM |
| 48 | R_inferiortemporal_1 | 45 | R_inferiortemporal_1 | LIM |
| 49 | R_inferiortemporal_2 | 46 | R_inferiortemporal_2 | DAN |
| 50 | R_middletemporal_1 | 47 | R_middletemporal_1 | DMN |
| 51 | R_middletemporal_2 | 48 | R_middletemporal_2 | DMN |
| 52 | R_bankssts_1 | 49 | R_bankssts_1 | SAL/VAN |
| 53 | R_superiortemporal_1 | 50 | R_superiortemporal_1 | SMN |
| 54 | R_superiortemporal_2 | 51 | R_superiortemporal_2 | DMN |
| 55 | R_transversetemporal_1 | 52 | R_transversetemporal_1 | SMN |
| 56 | R_insula_1 | 53 | R_insula_1 | SMN |
| 57 | R_insula_2 | 54 | R_insula_2 | SAL/VAN |
| 58 | R_thalamusproper | 55 | R_thalamusproper | SUB |
| 59 | R_caudate | 56 | R_caudate | SUB |
| 60 | R_putamen | 57 | R_putamen | SUB |
| 61 | R_pallidum | 58 | R_pallidum | SUB |
| 62 | R_accumbensarea | 59 | R_accumbensarea | SUB |
| 63 | R_hippocampus | 60 | R_hippocampus | LIM |
| 64 | R_amygdala | 61 | R_amygdala | LIM |
| 65 | L_lateralorbitofrontal_1 | 62 | L_lateralorbitofrontal_1 | LIM |
| 66 | L_lateralorbitofrontal_2 | 63 | L_lateralorbitofrontal_2 | LIM |
| 67 | L_parsorbitalis_1 | 64 | L_parsorbitalis_1 | DMN |
| 68 | L_frontalpole_1 | 65 | L_frontalpole_1 | LIM |
| 69 | L_medialorbitofrontal_1 | 66 | L_medialorbitofrontal_1 | DMN |
| 70 | L_parstriangularis_1 | 67 | L_parstriangularis_1 | DMN |
| 71 | L_parsopercularis_1 | 68 | L_parsopercularis_1 | SAL/VAN |
| 72 | L_rostralmiddlefrontal_1 | 69 | L_rostralmiddlefrontal_1 | CONT |

| | | | | |
|---|---|---|---|---|
| 73 | L_rostralmiddlefrontal_2 | 69 | L_rostralmiddlefrontal_2 | CONT |
| 74 | L_rostralmiddlefrontal_3 | 70 | L_rostralmiddlefrontal_3 | DMN |
| 75 | L_superiorfrontal_1 | 71 | L_superiorfrontal_1 | DMN |
| 76 | L_superiorfrontal_2 | 72 | L_superiorfrontal_2 | DMN |
| 77 | L_superiorfrontal_3 | 73 | L_superiorfrontal_3 | DMN |
| 78 | L_superiorfrontal_4 | 74 | L_superiorfrontal_4 | SAL/VAN |
| 79 | L_caudalmiddlefrontal_1 | 75 | L_caudalmiddlefrontal_1 | DAN |
| 80 | L_precentral_1 | 78 | L_precentral_1 | SMN |
| 81 | L_precentral_2 | 78 | L_precentral_2 | SMN |
| 82 | L_precentral_3 | 77 | L_precentral_3 | SMN |
| 83 | L_precentral_4 | 76 | L_precentral_4 | SAL/VAN |
| 84 | L_paracentral_1 | 79 | L_paracentral_1 | SMN |
| 85 | L_rostralanteriorcingulate_1 | 80 | L_rostralanteriorcingulate_1 | DMN |
| 86 | L_caudalanteriorcingulate_1 | 81 | L_caudalanteriorcingulate_1 | SAL/VAN |
| 87 | L_posteriorcingulate_1 | 82 | L_posteriorcingulate_1 | CONT |
| 88 | L_isthmuscingulate_1 | 83 | L_isthmuscingulate_1 | DMN |
| 89 | L_postcentral_1 | 85 | L_postcentral_1 | SMN |
| 90 | L_postcentral_2 | 85 | L_postcentral_2 | SMN |
| 91 | L_postcentral_3 | 84 | L_postcentral_3 | SMN |
| 92 | L_supramarginal_1 | 87 | L_supramarginal_1 | SAL/VAN |
| 93 | L_supramarginal_2 | 86 | L_supramarginal_2 | CONT |
| 94 | L_superiorparietal_1 | 88 | L_superiorparietal_1 | DAN |
| 95 | L_superiorparietal_2 | 89 | L_superiorparietal_2 | DAN |
| 96 | L_superiorparietal_3 | 90 | L_superiorparietal_3 | VIS |
| 97 | L_inferiorparietal_1 | 92 | L_inferiorparietal_1 | DAN |
| 98 | L_inferiorparietal_2 | 91 | L_inferiorparietal_2 | DMN |
| 99 | L_precuneus_1 | 94 | L_precuneus_1 | DMN |
| 100 | L_precuneus_2 | 93 | L_precuneus_2 | DMN |
| 101 | L_cuneus_1 | 95 | L_cuneus_1 | VIS |
| 102 | L_pericalcarine_1 | 96 | L_pericalcarine_1 | VIS |
| 103 | L_lateraloccipital_1 | 97 | L_lateraloccipital_1 | VIS |
| 104 | L_lateraloccipital_2 | 98 | L_lateraloccipital_2 | VIS |
| 105 | L_lingual_1 | 99 | L_lingual_1 | VIS |
| 106 | L_lingual_2 | 100 | L_lingual_2 | VIS |
| 107 | L_fusiform_1 | 101 | L_fusiform_1 | VIS |
| 108 | L_fusiform_2 | 102 | L_fusiform_2 | LIM |
| 109 | L_parahippocampal_1 | 103 | L_parahippocampal_1 | LIM |
| 110 | L_entorhinal_1 | 104 | L_entorhinal_1 | LIM |
| 111 | L_temporalpole_1 | 105 | L_temporalpole_1 | LIM |

| | | | | |
|---|---|---|---|---|
| 112 | L_inferiortemporal_1 | 106 | L_inferiortemporal_1 | LIM |
| 113 | L_inferiortemporal_2 | 107 | L_inferiortemporal_2 | DAN |
| 114 | L_middletemporal_1 | 108 | L_middletemporal_1 | DMN |
| 115 | L_middletemporal_2 | 109 | L_middletemporal_2 | DMN |
| 116 | L_bankssts_1 | 110 | L_bankssts_1 | DMN |
| 117 | L_superiortemporal_1 | 111 | L_superiortemporal_1 | SMN |
| 118 | L_superiortemporal_2 | 112 | L_superiortemporal_2 | DMN |
| 119 | L_transversetemporal_1 | 113 | L_transversetemporal_1 | SMN |
| 120 | L_insula_1 | 114 | L_insula_1 | SMN |
| 121 | L_insula_2 | 115 | L_insula_2 | SAL/VAN |
| 122 | L_thalamusproper | 116 | L_thalamusproper | SUB |
| 123 | L_caudate | 117 | L_caudate | SUB |
| 124 | L_putamen | 118 | L_putamen | SUB |
| 125 | L_pallidum | 119 | L_pallidum | SUB |
| 126 | L_accumbensarea | 120 | L_accumbensarea | SUB |
| 127 | L_hippocampus | 121 | L_hippocampus | LIM |
| 128 | L_amygdala | 122 | L_amygdala | LIM |
| 129 | brainstem | 123 | brainstem | Not assigned |

* The brainstem is excluded from the analysis due to insufficient coverage in some subjects. Abbreviations: VIS, visual network; SMN, somatomotor network; DAN, dorsal attention network; SAL/VAN, salience/ventral attention network; CONT, executive control network; DMN, default mode network; SUB, subcortical network.
